# Structural insights into the mechanism of C3 side chain fragmentation in cephalosporins with SME-1 class A carbapenemase

**DOI:** 10.1101/2025.07.17.665329

**Authors:** Kunal Dhankhar, Gourav Chakraborty, Abhishek Bera, Arnab Chowdhary, Anupam Bandyopadhyay, Narayan C Mishra, Niladri Patra, Saugata Hazra

## Abstract

Cephalosporins remain one of the largest classes of antibiotics used clinically, with a diverse group of dual substituents at C3 and C7 positions. The fragmentation of the C-3 side chain from the C3’ position after hydrolysis by Serine or metallo-β lactamase leaving an exo-methylene group has been introduced in the literature recently, but the exact mechnism by which it happens remains unclear until now. Our *incrystallo* findings suggest that the fragmentation of the C3 side chain will depend on the C3’ atom and its hybridization state. A C3’ carbon with sp2 hybridization will not allow fragmentation, but a sp3 hybridized carbon will facilitate the detachment of the side chain. Furthermore, heteroatoms like Sulfur, on C3’ position, bearing vacant d-orbitals participate in π-cloud delocalization, making the detachment of C3 side chain energetically unfeasible. For the sp3 hybridized carbon at C3’, the fragmentation of the bond was independent of the C3’’ atom. We observed fragmentation of the C3’-C3’’ bond for all three heteroatoms, Nitrogen, Oxygen, and sulfur, at the C3’’ position. We validated our crystallographic findings with mass spectrometry to confirm the fragmentation of the C3 side chain in sp3 carbon and the unfragmented side chain in sp2 carbon and sulfur at C3’ position. QM/MM simulations on three types of enzyme-cephalosporin complexes with different C3 side chains confirm the mechanism of fragmentation. Beyond revealing C3 side chain fragmentation in cephalosporins, useful for inhibitor and diagnostic design, our QM/MM methods are useful on *incrystallo* complexes to show the enzyme catalysis.

## Introduction

β-lactams are the largest and the most widely used class of antibiotics for treating bacterial infections. Since the discovery of penicillin in 1929, β-lactams have transformed modern medicine, making previously fatal infections curable. Among β-lactams, cephalosporins have played a pivotal due to their broad-spectrum activity and improved resistance to β-lactamases. These antibiotics are characterized by a 4-membered azetidin-2-one ring fused with a 6-membered dihydrothiazine moiety^1^. This azetidine-2-one ring lies at the core of all Cephalosporins and inhibits the activity of bacterial transpeptidases by blocking the crosslinking of NAG-NAM peptidoglycan units during cell-wall synthesis, ultimately causing bacterial cell death^2^. However, the widespread use of Cephalosporins has driven the evolution of bacterial resistance mechanisms, diminishing their efficacy. Resistance often arises via combination of factors, including target modifications (mutations or altered Penicillin Binding Proteins [PBPs]), reduced membrane permeability via porin downregulation, increased efflux activity, and production of β-lactamases^3^. The latter, in particular, poses a major challenge as these enzymes hydrolyze the β-lactam ring, rendering these antibiotics inactive. To counteract this, newer generation Cephalosporins have been developed with chemical modifications at the C3 and C7 side chains, enhancing their resistance to degradation. 4^th^ and 5^th^ generation Cephalosporins,for instance, feature bulky C7 oxyimino groups^4^, conferring stability against serine β-lactamases and activity against resistant PBPs, such as PBP2a from methicillin-resistance *Staphylococcus aureus*^5,6^. Despite their robustness, these newer cephalosporins remain prone to hydrolysis by Class A carbapenemases and Class B Metallo-β-lactamases (MBLs), including the clinically significant New Delhi Metallo-β-lactamase (NDM-1), further contributing to bacterial resistance^7,8^. Additionally, extended-spectrum-β-lactamases (ESBLs) such as FTU-1 can inactivate oxyimino-cephalosporins by targeting their C7 sidechains^9^.

Interestingly, recent studies have revealed that cephalosporins bearing specific C3 substitutions exhibit unique hydrolysis induced fragmentation behavior, which may have therapeutic implications^10^. Upon β-lactam ring hydrolysis, certain C3 sidechains spontaneously fragment, potentially releasing secondary bioactive compounds. This phenomenon, observed in cephalosporins with electron-acceptor groups at C3, can be useful for a dual-action antibacterial effect^11^. For instance, the cephalosporin MCO releases 2-mecaptopyridine-N-oxide, an antimicrobial compound, upon hydrolysis by β-lactamases, thereby enhancing its antibacterial activity^12^. The plausible mechanism involves an imine-enamine tautomerism, wherein the double bond migrates from C3=C4 (enamine) to C4=N5 (imine), with a concomitant transfer of proton to C3^13,14^. This protonated cephalosporoate intermediate can undergo different rearrangements, such as fragmentation of C3 side chain, yielding exo-methylene products or shifting of conjugated dienes, or no change at all.

In this study, we aimed to investigate the interaction of cephalosporins with Class A Carbapenemases, with a particular focus on SME-1. We determined the structure of SME-1’s deacylation-deficient mutant (E166A) in complex with cephalosporins bearing varied C3 sidechains. Using enzyme kinetics, isothermal titration calorimetry (ITC), and mass spectrometry, we characterized the hydrolysis rates and thermodynamic interactions^15^. Additionally, microbiological assays, including antimicrobial susceptibility testing, provided insights into the bactericidal efficacy of these agents. Our work, therefore, attempts to provide a deeper understanding of Cephalosporin interactions with Class A Carbapenemases, with a focus on C3’ behavior impacting the fragmentation of C3’-C3’’ bond.

## Results

### 4^th^ and 5^th^ generation cephalosporins are relatively resistant to hydrolysis by SME-1

All 5^th^-generation cephalosporins were poorer substrates to SME-1 with a catalytic efficiency of around 0.3 µM^−1^min^−1^. The enzyme’s catalytic efficiency was much poorer than even carbapenems, making them very potent compounds against this SME-1 Carbapenemase. With Cefaclor and Cefoperazone, the enzyme had a very good catalytic efficiency for hydrolysis, with values of 24.97 µM^−1^min^−1^ for cefaclor and 9.07 µM^−1^min^−1^ for Cefoperazone (Table 1). Cefaclor, being a very small compound, might be able to fit better during the formation of the acyl-enzyme complex, and a small molecule should be a better-leaving compound than a bulky compound like Cefoperazone. Being very bulky, other cephalosporins are not a good fit in SME-1’s active site. The catalytic efficiency was the poorest with Cefsulodin, which can be attributed to the presence of the sulfate group in the C7 side chain. Cefotetan, with a unique C7 side chain, was also a poorer substrate with a catalytic efficiency of 0.11 µM^−1^min^−1^, which was similar to that of 4th and 5th-generation cephalosporins (Table 1).

**Table 1.** Enzyme kinetics parameters for cephalosporins against SME-1.

| S.No. | Cephalosporin | $V_{\max}$ ($\mu\text{M}/\text{min}$) | $K_m$ ($\mu\text{M}$) | $k_{\text{cat}}$ ($\text{min}^{-1}$) | $k_{\text{cat}}/K_m$ ($\mu\text{M}^{-1} \text{min}^{-1}$) |
| --- | --- | --- | --- | --- | --- |
| 1 | Cefaclor | 385.18±14.81 | 3206.8±570 | 77037±2962 | 24.97±5.37 |
| 2 | Cefotetan | 12.04± 6.16 | 579.23±346 | 60.24±30.83 | 0.11±0.013 |
| 3 | Cefoperazone | 19.52 ±3.1 | 149.9±37.43 | 1328±211.03 | 9.07±0.85 |
| 4 | Cefsulodin | 12.55 ±2.73 | 643.5 ±124.3 | 62.78 ±12.66 | 0.097 ±0.0024 |
| 5 | Cefpirome | 16.47±2.68 | 180.75±2.13 | 82.37±14.3 | 0.45±0.0084 |
| 6 | Ceftaroline<br>Fosamil | 19.17 ±1.19 | 346.8 ±59 | 95.88 ±5.92 | 0.281 ±0.03 |
| 7 | Ceftobiprole | 215.55 ±78.56 | 3355.77<br>±1438 | 1077.7 ±392 | 0.331 ±0.0251 |

Cephalosporins with different side chains were studied for their binding with the help of Isothermal titration calorimetry with SME-1 deacylation-deficient mutant. All the cephalosporins showed a value for N around 1, which indicates that even after the rearrangement of the C3 side chain, it has no effect on the stoichiometry of ligand and enzyme. The Gibbs free energy for the system was lowest in the case of Ceftaroline with a value of −3.87 kcal/mol, indicating the poorest binding, which can be expected as it has the bulkiest side chain out of all used cephalosporins (Table 2). This C3 side chain does not leave after the formation of the acyl-enzyme adduct, which further creates steric hindrance with the enzyme, making the binding much poorer than the rest. Cefsulodin and Cefoperazone showed Gibbs energy −6.44 and −6.94 kcal/mol, respectively (Table 2). It indicates the fit of these two cephalosporins is much better than the Ceftaroline, as their C3 side chain has the potential to get removed. Cefpirome, being the 4th generation cephalosporin and also having a detachable C3 side chain, also bound well with the enzyme with a binding energy of −21.7 kcal/mol, which is more than the rest of the cephalosporins. Cefaclor, being the most easily cleaved cephalosporin, fits well in the active site as it only has an exocyclic halogen at the C3 side chain; given that it is one of the smallest cephalosporins, it has a Gibbs energy value of −37.7 kcal/mol (Table 2). Lastly, ceftobiprole, having a conjugated diene in its structure that shifts to form an imine and an exocyclic methylene group, showed the highest value for Gibbs’s energy of −76.9. This compound also showed a uniquely high value for −TdS. The dissociation constant KD for all these compounds was about in the micromolar range, with the lowest value for Cefoperazone and the highest value for ceftobiprole and cefaclor (Figure 2).

**Table 2.**
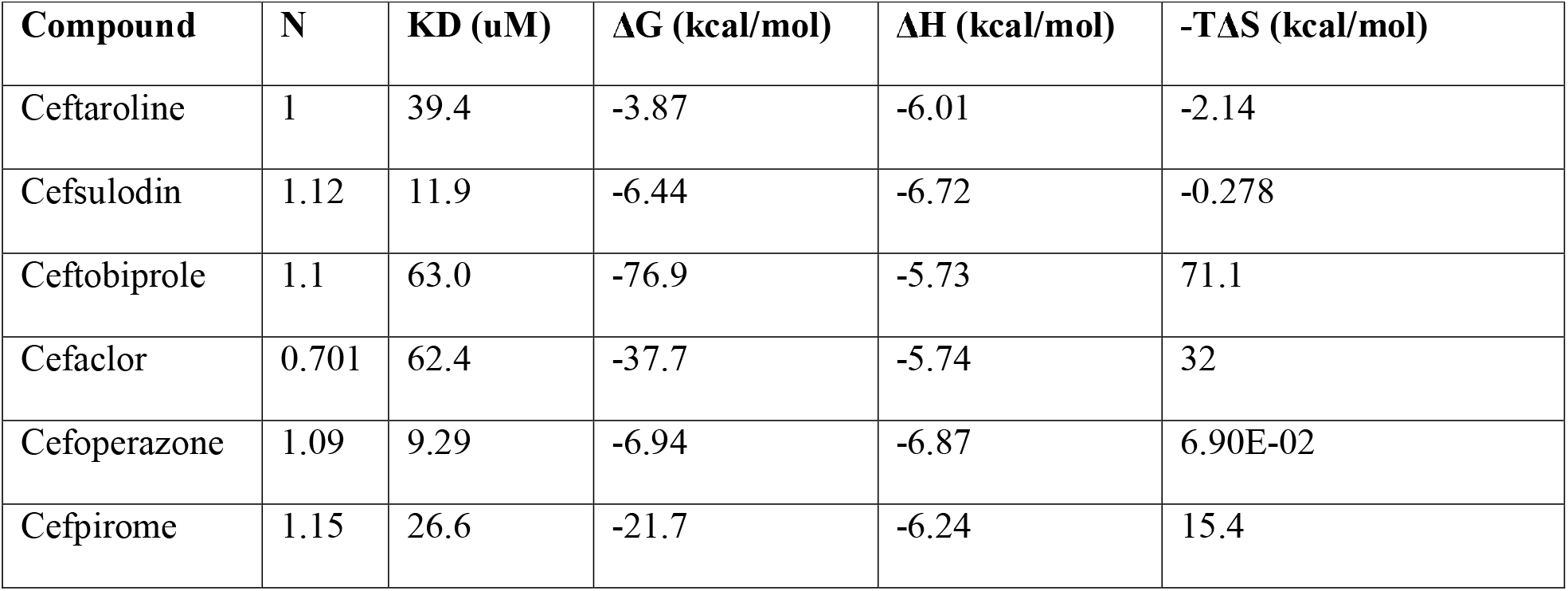
Binding parameters for cephalosporins against SME-1 E166A deacylation deficient mutant.

**Figure 1.**
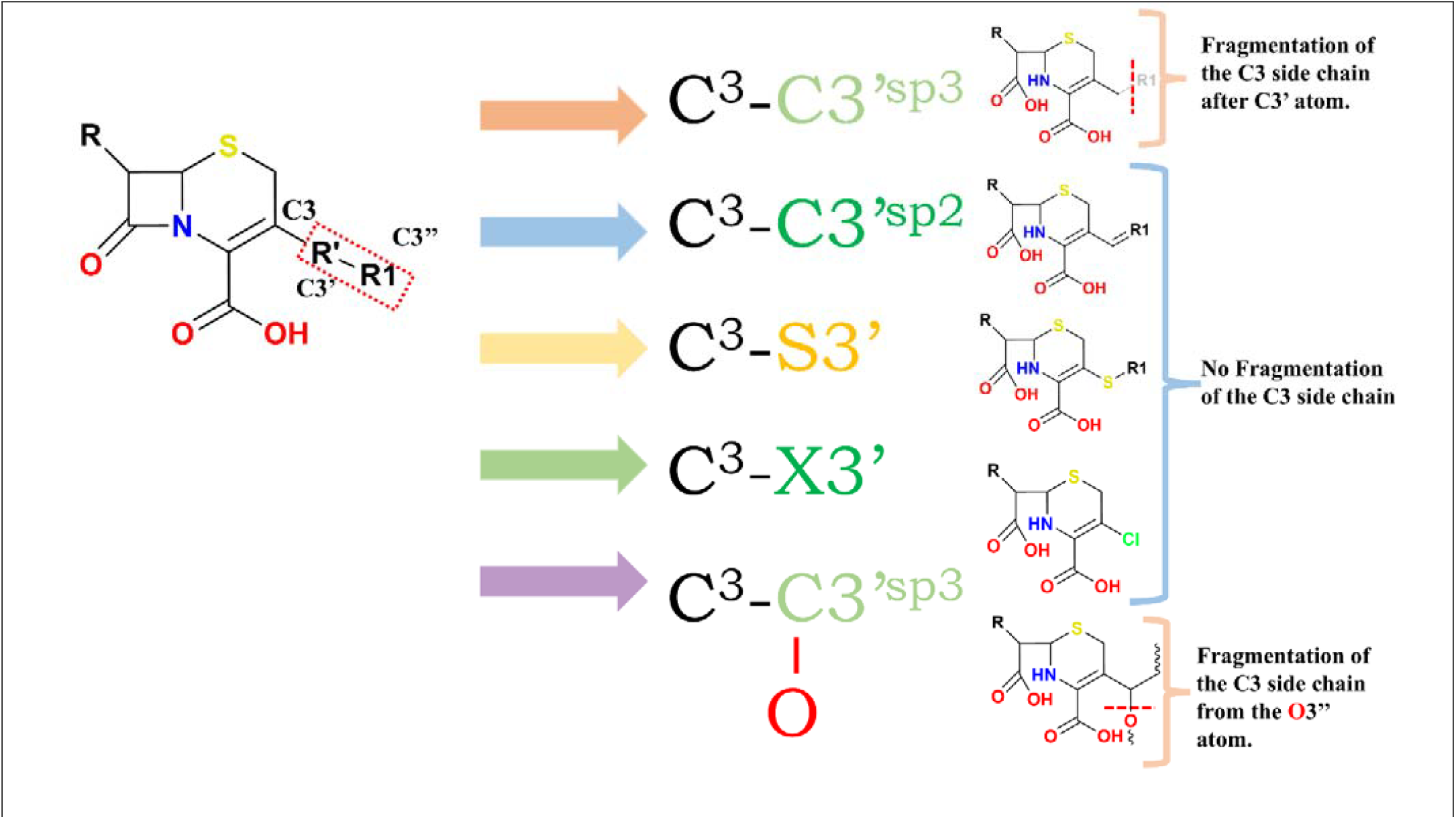
Cephalosporins with different types of side chains at the C3 position.

**Figure 2.**
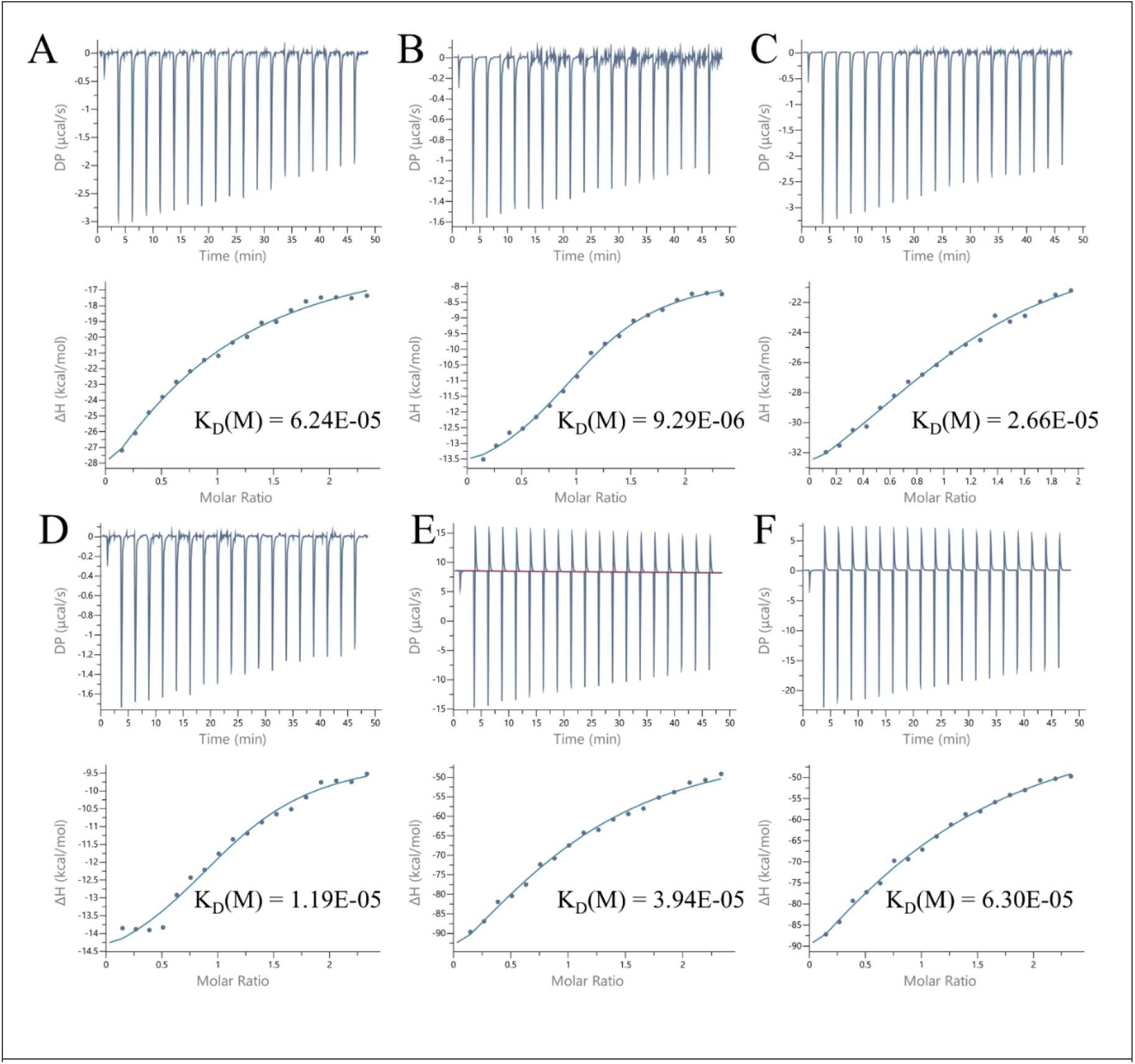
Isothermal titration calorimetry profile of Cephalosporins. (A) Cefaclor (B) Cefoperazone (C) Cefpirome (D) Cefsulodin (E) Ceftaroline Fosamil (F) Ceftobiprole

### Mass spectrometry of SME-1 deacylation-deficient mutant with cephalosporins

The mass of SME-1 E166A protein was 30124-30132 (Figure 3), and it was measured from two different batches of purified protein. Cefoperazone with an sp3 hybridized carbon atom in the C3 position showed the side chain fragmentation as the adduct mass was increased by 530 Daltons instead of about 645. This mass difference of 530 Daltons complements the formation of an exocyclic methylene product (Figure 3). Ceftobiprole with an sp2 hybridized C3’ carbon showed no signs of fragmentation of the C3 side chain in the acyl-enzyme complex. A mass difference of 534 Daltons was observed between the acyl-enzyme complex and the apoenzyme instead of 383 Daltons if an exomethylene product had been formed. With Ceftaroline Fosamil, which has a C3’ Sulphur instead of carbon, a mass difference of 684 Daltons was observed, suggesting that the presence of heteroatom at C3’ completely stops the fragmentation of the C3 side. Mass spectrometry with cefixime also revealed the same result as in the case of ceftobiprole (Figure 3).

**Figure 3.**
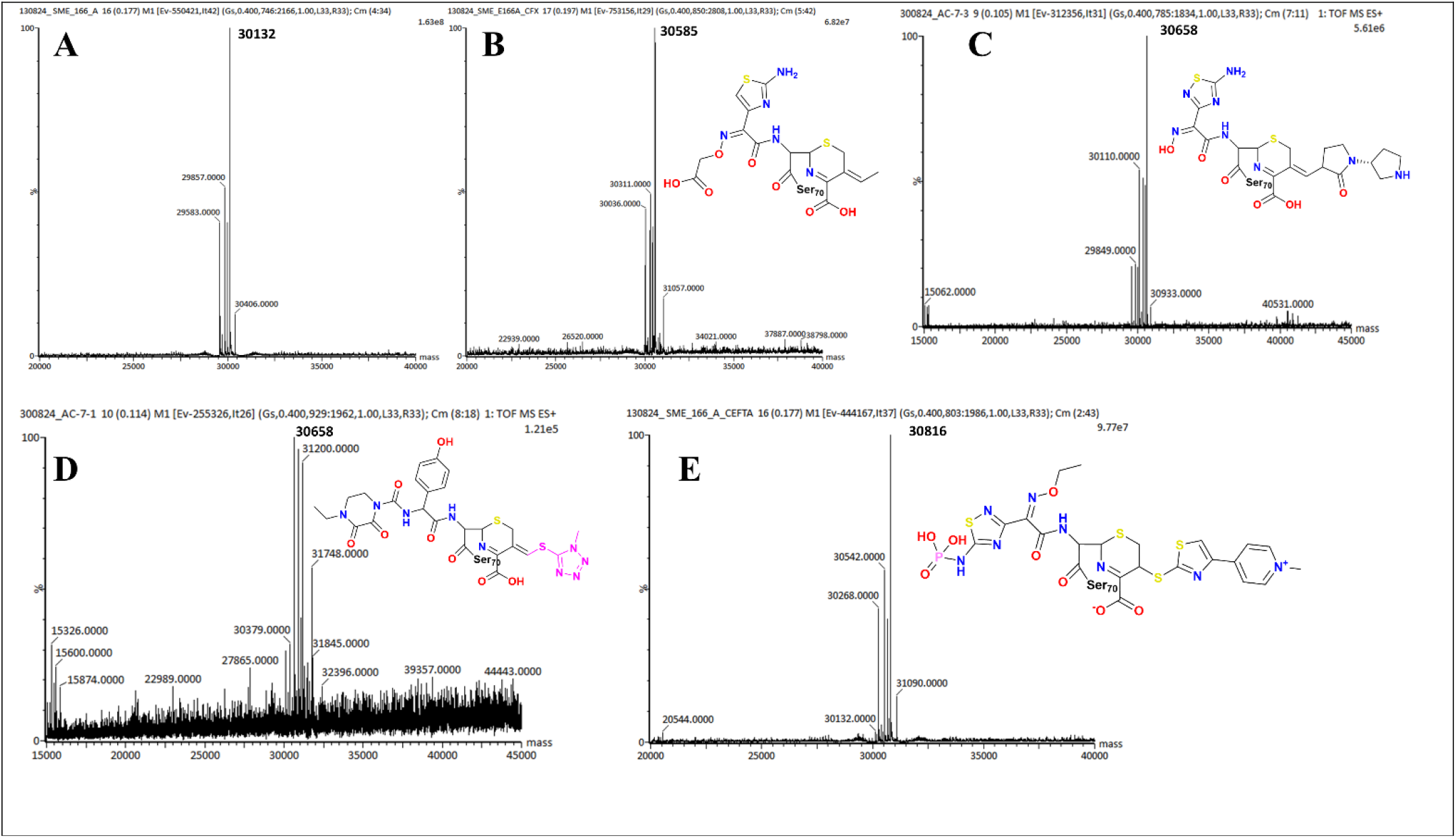
Mass spectra of SME-1 E166A with cephalosporins in acyl-enzyme complex. (A) Mass spectra of SME-1 E166A. (B) Mass spectra of SME-1 E166A with Cefixime. (C) Mass spectra of SME-1 E166A with Ceftobiprole. (D) Mass spectra of SME-1 E166A with Cefoperazone (C3 side chain fragmented, highlighted in pink). (E) Mass spectra of SME-1 E166A with Ceftaroline Fosamil.

### X-ray crystallography reveals the requirements for C3 side chain fragmentation in cephalosporins

Cefaclor, being the 1st generation cephalosporin and the smallest one used here, has only an exocyclic halogen at the C3 position. Formation of Δ1 Imine should be spontaneous in this case as a heteroatom has been present at the C3 position. As chlorine generally forms only 1 covalent bond, there is no C3’’ atom present from which we can study the fragmentation pattern of the same; chlorine being able only to form 1 bond does not show the presence of a double bond as seen in exomethylene adducts. The behaviour of the cefaclor with the C3 side chain can be compared with the carbapenems, as they also don’t show any C-side chain fragmentation. The carboxylic acid group shows the conserved interaction with the KTG motif, which is essential for binding beta-lactams. Ser130 forms an H-bond with the Imine nitrogen present in the ring. In SME-1, there is a serine at the 237 position, which essentially stabilizes the carbonyl oxygen with an H-bond (Figure 4 & 7).

**Figure 4.**
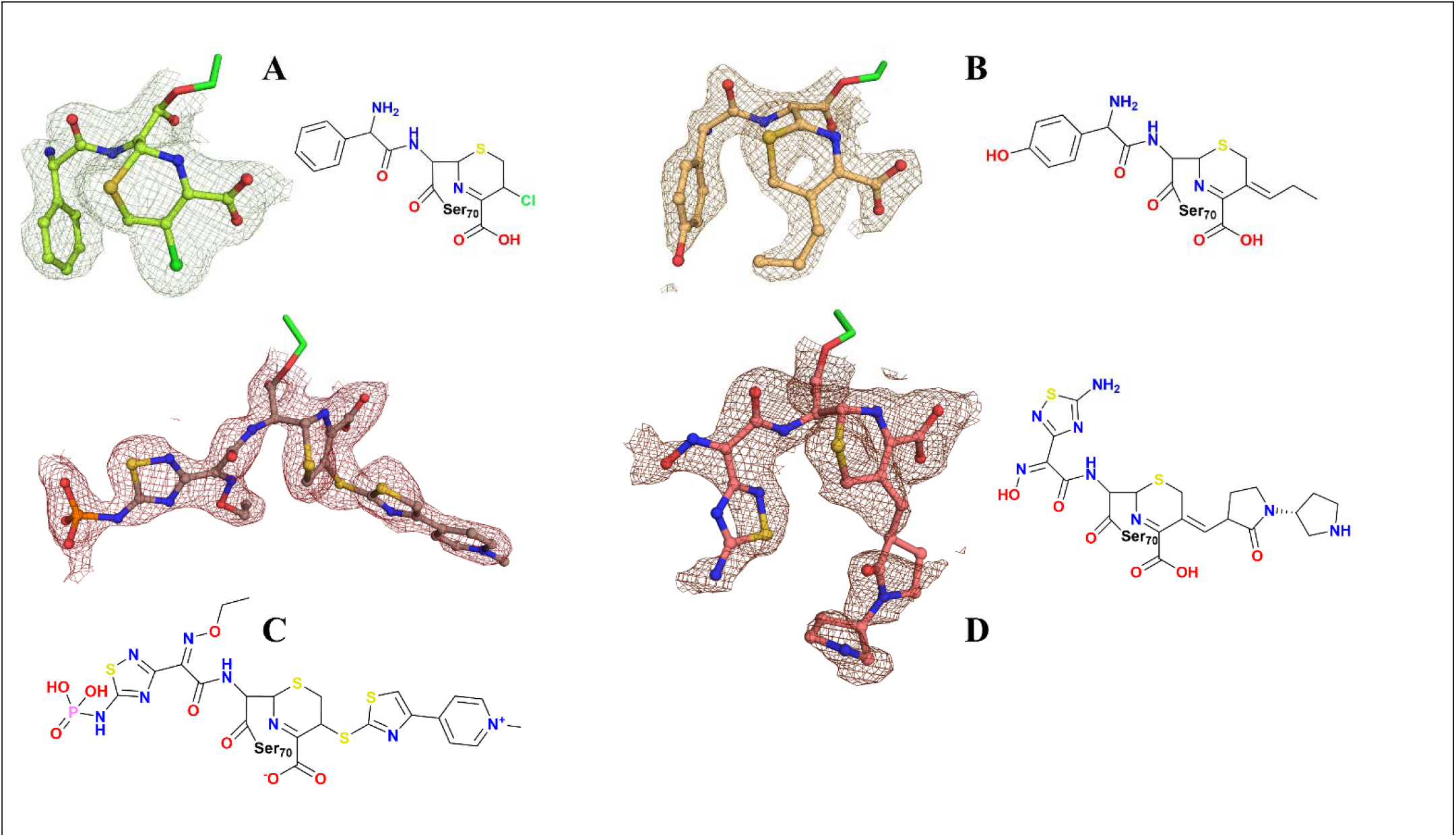
Crystal structures of cephalosporins retaining C3 side chains after acylation with SME-1 E166A. (A) The structure of Cefaclor showing a halogen substituent does not allow the formation of an exomethylene adduct. (B) The structure of Cefprozil with a sp2 hybridized C3’ atom shows rearrangement of the pi cloud and retaining of the C3 side chain. (C) The structure of Ceftaroline with sulfur does not allow the formation of the exomethylene adduct and also retains the C3 side chain. (D) Structure of Ceftobiprole with a sp2 hybridized C3’ atom shows rearrangement of the pi cloud and retaining of the C3 side chain.

Cefotetan has a unique oxy-methyl group at the C7 position, which forms a critical H-bond with lys73 of the enzyme. This might explain the lower catalytic efficiency of SME-1 against cefotetan. Imine nitrogen of beta-lactam ring forms H-bond with Ser130 of SDN Loop. The carboxylic acid group interacts with the KTG motif of the SME-1, which is conserved throughout the beta-lactams (Figure 5 & 7).

**Figure 5.**
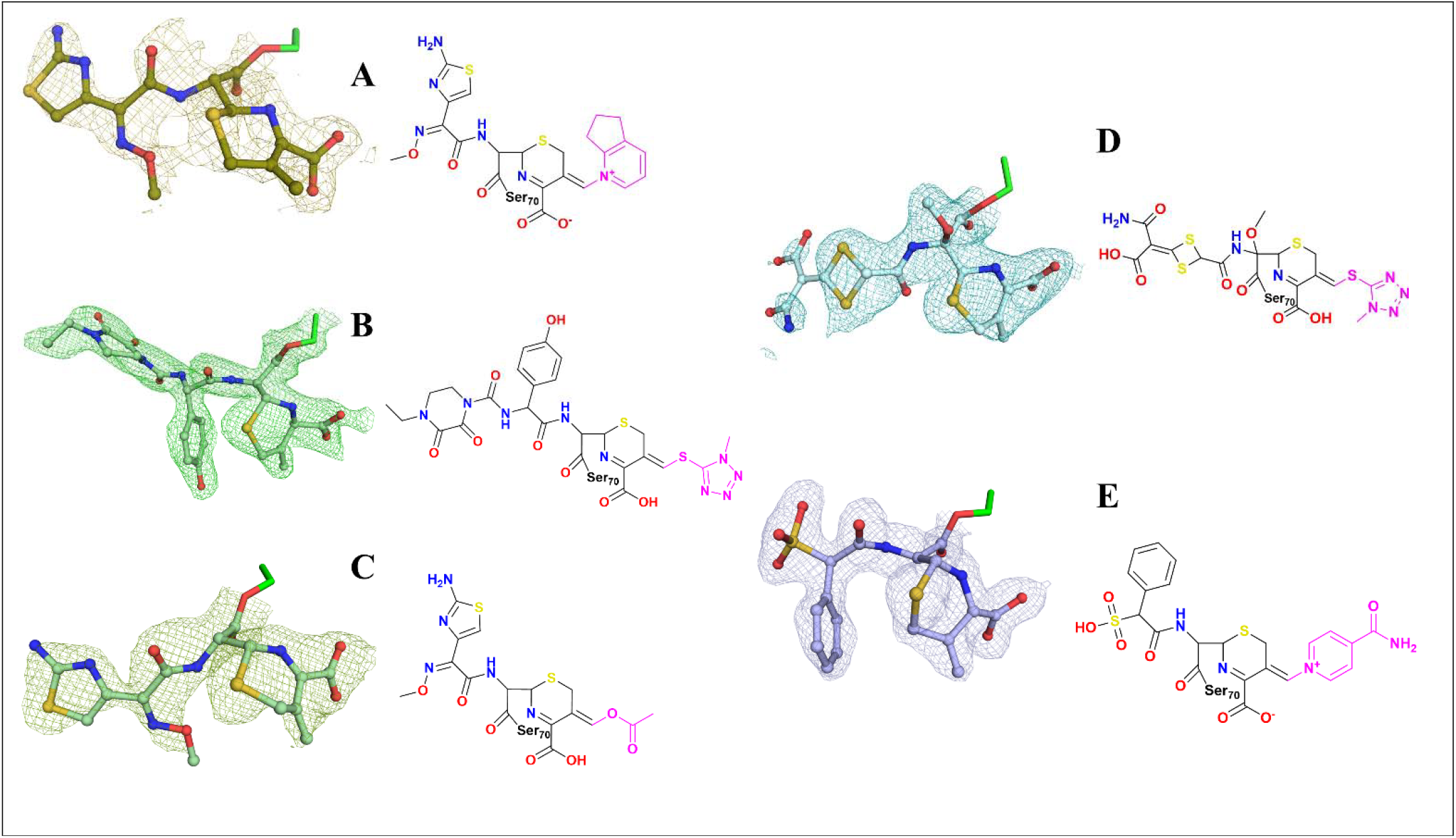
Crystal structures of cephalosporins with fragmented C3 side chains after acylation with SME-1 E166A (fragmented portions are shown in pink), leaving an exomethylene group. This fragmentation is independent of the atom from leaving group that is directly attached to the C3’ atom. (A) Structure of Cefpirome. (B) Structure of Cefoperazone. (C) Structure of Cefotaxime. (D) Structure of Cefotetan. (E) Structure of Cefsulodin.

Cefotaxime, being the oxyimino cephalosporin, forms additional H-bonds with the Asn132 from the SDN motif, Ser237, which is located just after the KTG motif, and also stabilizes the structure by some Hydrophobic interactions. Ser237 also interacts with the carboxylic acid group of the antibiotic. Lys234 forms ionic interactions with this carboxylic acid (Figure 5 & 7).

Cefovecin has a unique methine carbon atom at C3’ position because of the tetra hydro furan ring. This leads to two possibilities of fragmentation of the THF ring. Either the C-O or C-C bond can fragment, but because of the electronegativity of the oxygen and the polar nature of the C-O bond, it is more likely to undergo fragmentation. So, post acylation, the proton should transfer from C3 carbon to the O3’’ atom, resulting in the opening of the ring. In the acyl-enzyme complex of cefovecin, the carboxylic acid group forms H-bonds with the KTG motif and Ser130 of the beta-lactamase and an ionic interaction with the Lys234. The terminal hydroxyl of the C3 side chain forms an H-bond with His105. The primary amine substituent of the C7 thiazole substituent interacts with Leu167 from the E^166^XXXN^170^ (Ambler numbering) motif, providing extra stability to the adduct (Figure 6 & 7). Our H1 and C13 findings also support the crystallographic data where the C-O bond of the tetrahydrofuran (THF) has been fragmented. Although in the hydrolysed cefovecin, a mixture of both the unreacted cefovecin and fragmented cefovecin can be seen in the NMR, the shift of proton (d) (Figure S4 and S6) and carbon (d) (Figure S5 & S7) up field in the H1 & C13 NMR confirms the fragmentation of the C-O bond the THF sidechain.

**Figure 6.**
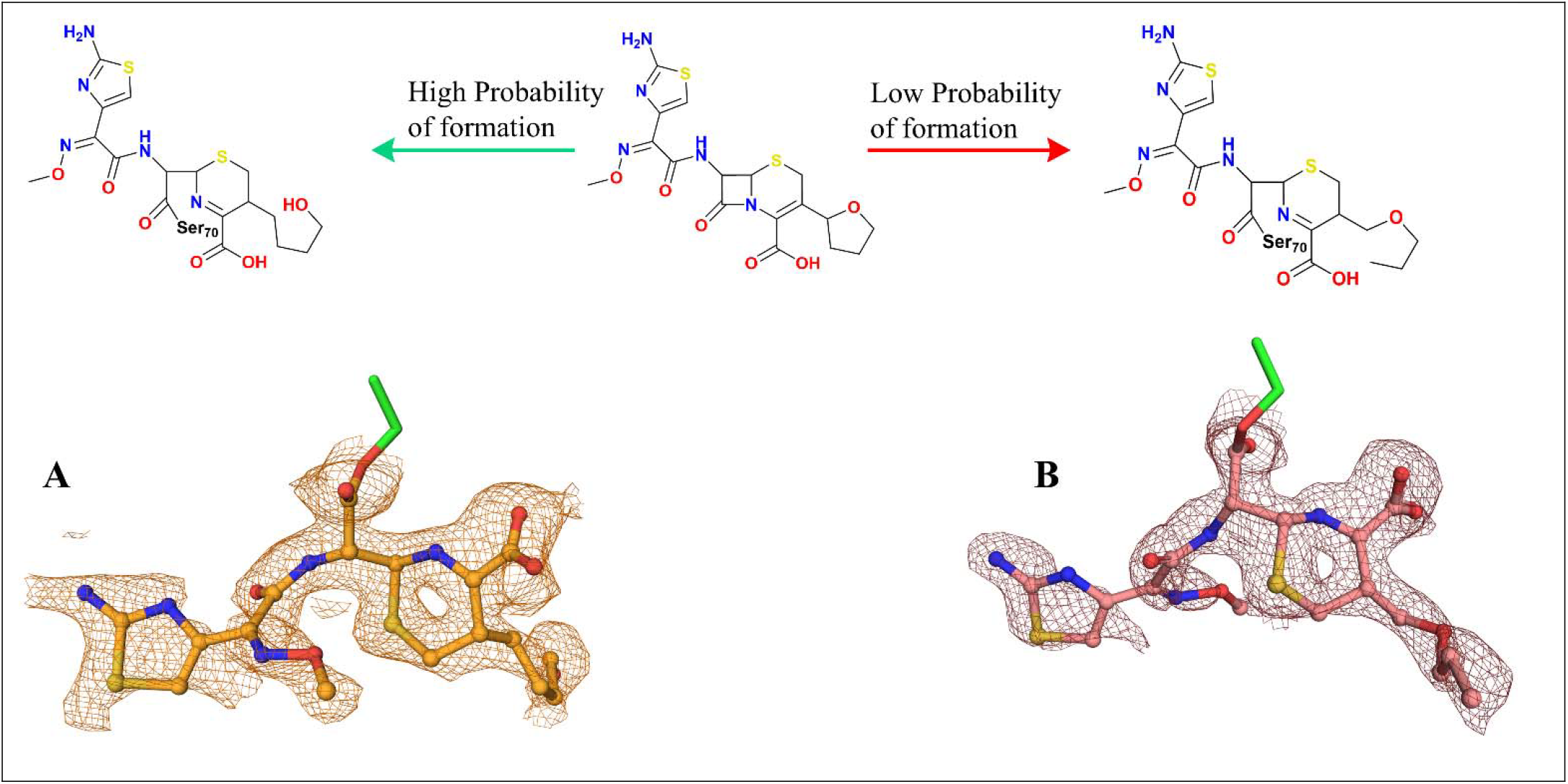
Crystal Structure of Cefovecin with SME-1 E166A. Arrows show two plausible pathways for fragmentation after hydrolysis of the beta-lactam ring. (A) C-O bind fragments, leaving an aliphatic alcohol side chain. (B) C-C bond fragments leaving an aliphatic ether. (Note: Both structural isomers were fitted in the electron density, and (A) was finalized as the final product that has formed after considering NMR Data.)

**Figure 7.**
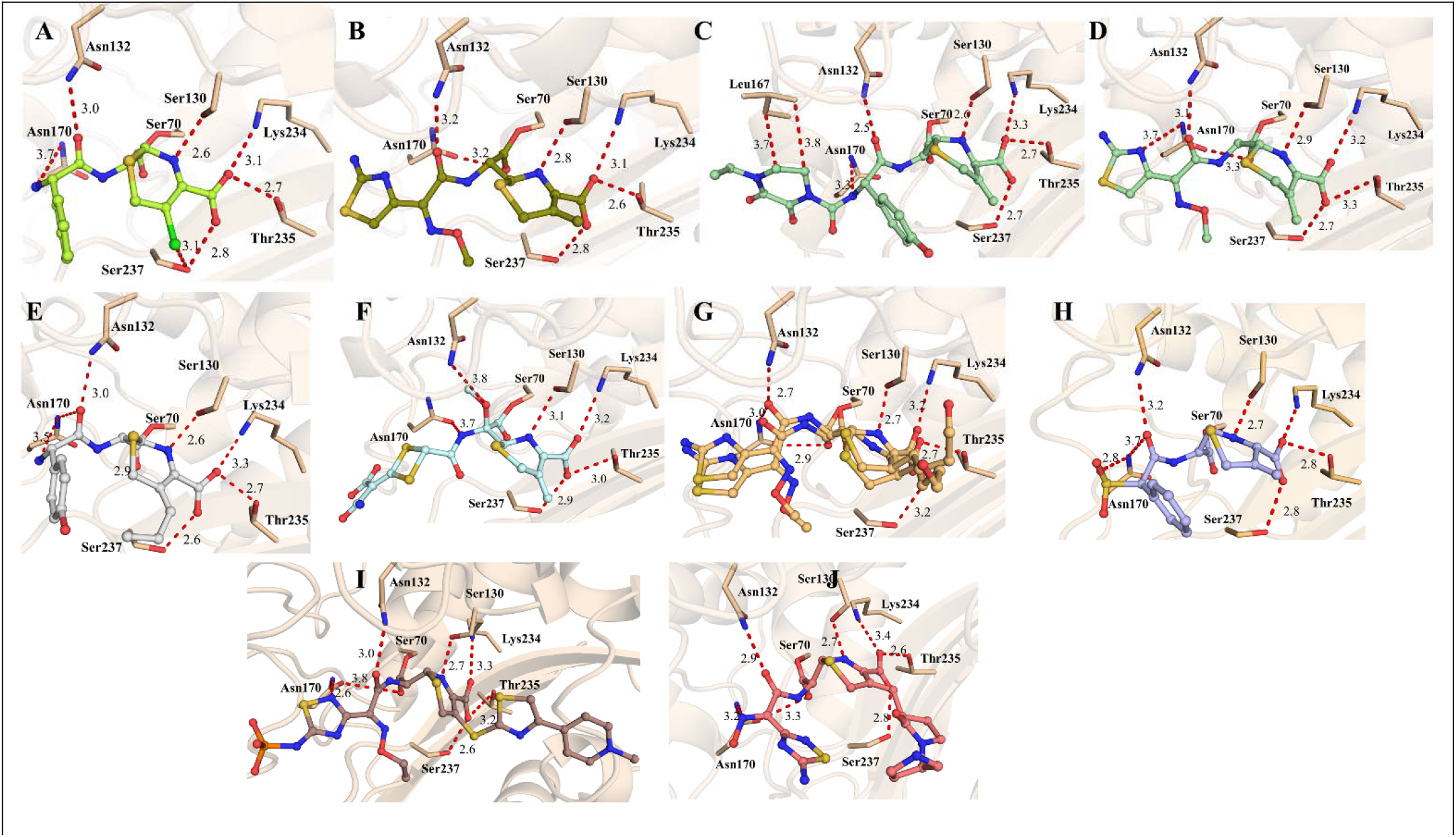
Interaction profile of cephalosporins with SME-1 E166A. (A) Cefaclor (B) Cefpirome (C) Cefoperazone (D) Cefotaxime (E) Cefprozil (F) Cefotetan (G) Cefovecin (H) Cefsulodin (I) Ceftaroline Fosamil(J) Ceftobiprole

Cefoperazone, being a very bulkier 3rd generation cephalosporin, binds the SME-1 E166A with some more interactions than the cefaclor. 1^st^ notable thing we can understand from its structure is that the fragmentation of the C3 side chain has taken place immediately after acylation. The exomethyl group has been converted to the exomethylene group, with the fragmentation taking place from the C3’’ sulfur atom. The whole tetrapyrrole side chain starting from sulfur got fragmented from the Cefoperazone, indicating that the C3’’ atom does not play any role in this spontaneous rearrangement. Only the atom at C3’s position determines whether the fragmentation will occur or not. The carboxylic group in Cefoperazone also showed a similar bind to cefaclor with mainly the KTG motif. Ser130 forms a H-bond with the imine group. The C7 side chain has some unique interactions like with Cys238 and Asn132 (Figure 5 & 7).

Cefsulodin has a similar sp3 hybridized carbon at the C3’ position like Cefoperazone, but it is followed by a nitrogen atom at the C3’’ position, unlike Cefoperazone, which has a sulfur. But immediately after the formation of the acyl-enzyme adduct, the whole C3 side chain after the C3’ carbon atom fragments from the molecule, forming the exocyclic methylene group; this leads to the formation of a conjugated diene in the structure of the cephalosporins. The carboxylic acid group forms a very similar interaction with the KTG motif like other cephalosporins; Arg220 also forms a long-range ionic interaction with this carboxylic acid group. One peculiar interaction of C7 side chain sulphate is with the Asn170, which is not found in any other cephalosporins. This interaction might explain the very less catalytic efficiency of the Cefsulodin by SME-1 Carbapenemase (Figure 5 & 7).

Like Cefsulodin and Cefoperazone, cefpirome also forms an exomethylene product after acylation with the enzyme. Being a 4^th^ generation cephalosporin, it has a very poor value for catalytic efficiency with SME-1, which can be explained by the additional interaction in the crystal structure. The carboxylic acid showed a conserved interaction with the KTG motif. The Asn132 forms an H-bond with the C7 side chain, and the Leu167 from the Omega loop forms an H-bond with the terminal amine group in the C7 side chain (Figure 5 & 7).

We selected Cefprozil as it has a C3 carbon atom but in the sp2 hybridization instead of sp3. It was observed that the atom might be like that of previous cephalosporins, but due to this change in hybridization, the whole C3 side chain remains intact, unlike the Cefoperazone, cefpirome, and Cefsulodin. The presence of this diene in the structure might have a significant impact on this fragmentation phenomenon. This π cloud shifts to form the same identical exomethylene group as in other sp3 hybridized cephalosporins. But these C3 side chains remain intact in this case. The rest of the interactions were very similar to those of other cephalosporins. The carboxylic side chain forms H-bonds with the KTG motif, and the amine group from the C7 side chain forms H-bonds with Ser237 (Figure 4 & 7).

Ceftobiprole having a sp2 hybridized atom at C3’ position with a bulky rest of the side chain motivated us to soak it with the SME-1 deacylation deficient mutant. If present, the electron density of the side chain can help us establish the exact role of the C3 atom in the fragmentation of the C3 side chain. Like Cefprozil, ceftobiprole’s side chain was intact upon acylation, indicating that the hybridization of the C3 atom also plays an important role in the fragmentation. This sp2 hybridized carbon atom avoids side chain fragmentation as the carbon-carbon double bond is harder to break than a single bond. However, this pi cloud shifted to form the imine isomer and the exocyclic methylene group, but unlike the sp3 carbon at C3’, the side chain is not fragmented in this case. This unfragmented C3 side chain contributes to additional hydrophobic interactions with a His273 and Ser237 atom, which stabilizes this complex more than the rest of lower generation cephalosporins. The rest of the interactions were very similar to those of other cephalosporins (Figure 4 & 7).

Ceftaroline Fosamil has a unique C3’ exocyclic sulfur atom, making it the only cephalosporin with this type of side chain. In its complex with the SME-1 E166A, there is a very clear electron density for the C3 side chain, which indicates that C3’ exocyclic sulfur also prevents the fragmentation of the side chain. This nature of sulfur can also be seen in the carbapenems as they also have a C2’ exocyclic structure, and their side chain can also be found in the acyl-enzyme complex. This side chain does the hydrophobic interactions with threonine at 216 and the histidine present at 104. This hydrophobicity might be the cause of the lower catalytic efficiency of the enzyme against this drug. The rest of the interaction of this compound with the KTG motif is similar to those of previous ones, with the C7 side chain making an additional H-bond with Asn170 (Figure 4 & 7).

### The C3’ atom alone defines the fragmentation of the C3 Sidechain

The nature of the C3’ atom determines how the C3 side chain in cephalosporins fragments. As seen with Cefoperazone, Cefsulodin, and rest carbon-containing cephalosporins, where the exo-methyl group is changed into an exomethylene (Figure 5), side chain fragmentation is caused due to sp3-hybridized C3’ carbons right after acylation. On the other hand, because of the stability of the C=C double bond (Figure 4) and the proton transfer, cephalosporins with sp2-hybridized C3’ carbons, like Cefprozil and ceftobiprole, maintain their side chains. Similar to carbapenems, Ceftaroline Fosamil, which has an exocyclic sulfur at C3’, likewise resists fragmentation, most likely as a result of the influence of the sulfur’s empty d-orbital. The binding interactions within the SME-1 active site are influenced by these structural differences.

Our findings suggest that the C3’ atom’s nature alone determines the C3 side chain’s fragmentation in cephalosporins; the nature of the next C3’’ atom has no effect on this process. The C3’’ atom is carbon in each of the cephalosporins (Figure 5), but its nature varies—it can be nitrogen (cefpirome and cefsulodin), oxygen (cefotaxime), or sulfur (cefoperazone and cefotetan). Notwithstanding these variations of different atoms at C3’’, fragmentation always happens when the C3’ atom occurs sp^3^-hybridization, supporting the idea that fragmentation is mostly influenced by the C3 atom’s hybridization state rather than the chemical makeup of C3’’. Even though the nature of the C3’’ atom varies, this observation is clear in structures where the side chain is lost just after acylation. Further evidence that the C3 atom itself is the deciding factor comes from the fact that neither sulfur nor nitrogen or oxygen at C3’’ can stop fragmentation. These results are consistent with earlier structures, highlighting the fact that sp^2^-hybridized C3 atoms keep their side chains while sp^3^-hybridized C3 carbons undergo spontaneous rearrangement and side chain loss. Therefore, regardless of the chemical characteristics of the nearby C3’’ atom, the fragmentation mechanism seems to be inherently connected to the nature and hybridization of the C3 carbon.

### Antimicrobial Susceptibility Testing with *E*.*coli* Top 10 cells harbouring SME-1 reveals the effect of different C3 side chains in cephalosporins

In the disk diffusion assay, SME-1 in TOP 10 cells showed resistance to 1^st^ and 3rd-generation cephalosporins, but 5th-generation cephalosporins were still showing very good bactericidal activity against SME-1-producing TOP 10 cells. The zone of inhibition clearly indicates the susceptibility of SME-1 to Ceftaroline Fosamil and Ceftobiprole, which confirms that the C3 side chain of these antibiotics slows the beta-lactam ring hydrolysis, which additional binding that, in turn, inhibits more PBPs, causing bacterial cell death. A similar trend was observed in broth when MIC was determined with these agents (Table 3).

**Table 3.** Antimicrobial susceptibility testing of cephalosporins against.

|  |  |  |  |  |  |
| --- | --- | --- | --- | --- | --- |
|  |  | **MIC in mcg/ml** |  | **Disk Diffusion Assay (mm)** |  |
| **S.No.** | **Compound** | **E. coli** | **E. coli Top 10 with SME-1** | **E. coli** | **E. coli Top 10 with SME-1** |
| **1** | Cefaclor | 1 | 4 | 18 | No Zone |
| **2** | Cefixime | 0.25 | 1 | 22 | 18 |
| **3** | Cefoperazone | 0.125 | 0.5 | 28 | 18 |
| **4** | Cefpirome | 0.25 | 0.5 | 26 | 24 |
| **5** | Ceftobiprole | 0.03 | 0.04 | 28 | 25 |
| **6** | Ceftaroline Fosamil | 0.03 | 0.03 | 28 | 24 |

### QM/MM simulations reveal the mechanism for post-acylation C3 side chain fragmentation in cephalosporins

The PMF values obtained from Step I and Step II for all the ligands are shown in Figure 8 (B). As observed, the highest energy barrier (in terms of PMF) in Step I exits for Ceftobiprole (83.76 kcal/mol), followed by Cefsulodin (75.09 kcal/mol) and Ceftaroline (72.10 kcal/mol). For Step II, this ordering undergoes a dynamic shift, with Ceftaroline exhibiting the highest PMF (58.25 kcal/mol), followed by Ceftaroline (45.02 kcal/mol) and Cefsulodin (38.20 kcal/mol) (Figure 8(B)). Henc, the first step shows a smaller relative change in PMF values across different ligands compared to Step II. Considering Ceftaroline as the baseline for Step I, the relative change in PMF for Cefsulodin and Ceftobiprole are 2.99 kcal/mol and 1.66 kcal/mol. On the other hand, in Step II, the lowest PMF is exhibited by Cefsulodin, with respect to which the energy changes observed for Ceftobiprole and Ceftaroline are 6.82 kcal/mol and 20.05 kcal/mol. These observations suggest that the second step could be the rate-limiting step in this two-step process. This can explain the C3 side detachment observed for Cefsulodin but not with Ceftaroline. One interesting point to note here is the lower PMF value observed for Ceftobiprole in Step II, although no detachment was observed. To explain this behaviour, the variation in distances observed between atom C3”/N3”-H, and C3’/S3’-C3”/N3”was plotted, as shown in Figure 9. In all the plots of Figure 9, the violet lines indicate the migration of H-atom, from C3 to C3” (for Ceftaroline and Ceftobiprole), or C3 to N3” (for Cefsulodin). During this migration, a significant increment in the C3’-N3’’/C3” distance is observed for Cefsulodin/Ceftaroline (Figure 9 (A), (B)). This indicates the detachment of the C3 side chain. For Ceftobiprole, however, the change in the C3’-C3” distance was very marginal, going from 1.3 to 1.5 Å, explaining the transition from a double bond to a single bond. Hence, no C3 side chain detachment was observed for Ceftobiprole. Hence, combining the results from Figure 8 and Figure 9, it is established that the C3 side chain gets dissociated most easily for Cefsulodin, and although, in theory, Ceftaroline can also exhibit this behaviour, the associated energy barrier for this process is very high. For Ceftobiprole, however, no side chain detachment is possible due to the existence of an initial double bond between C3’ and C3” (Figure 9).

**Figure 8:**
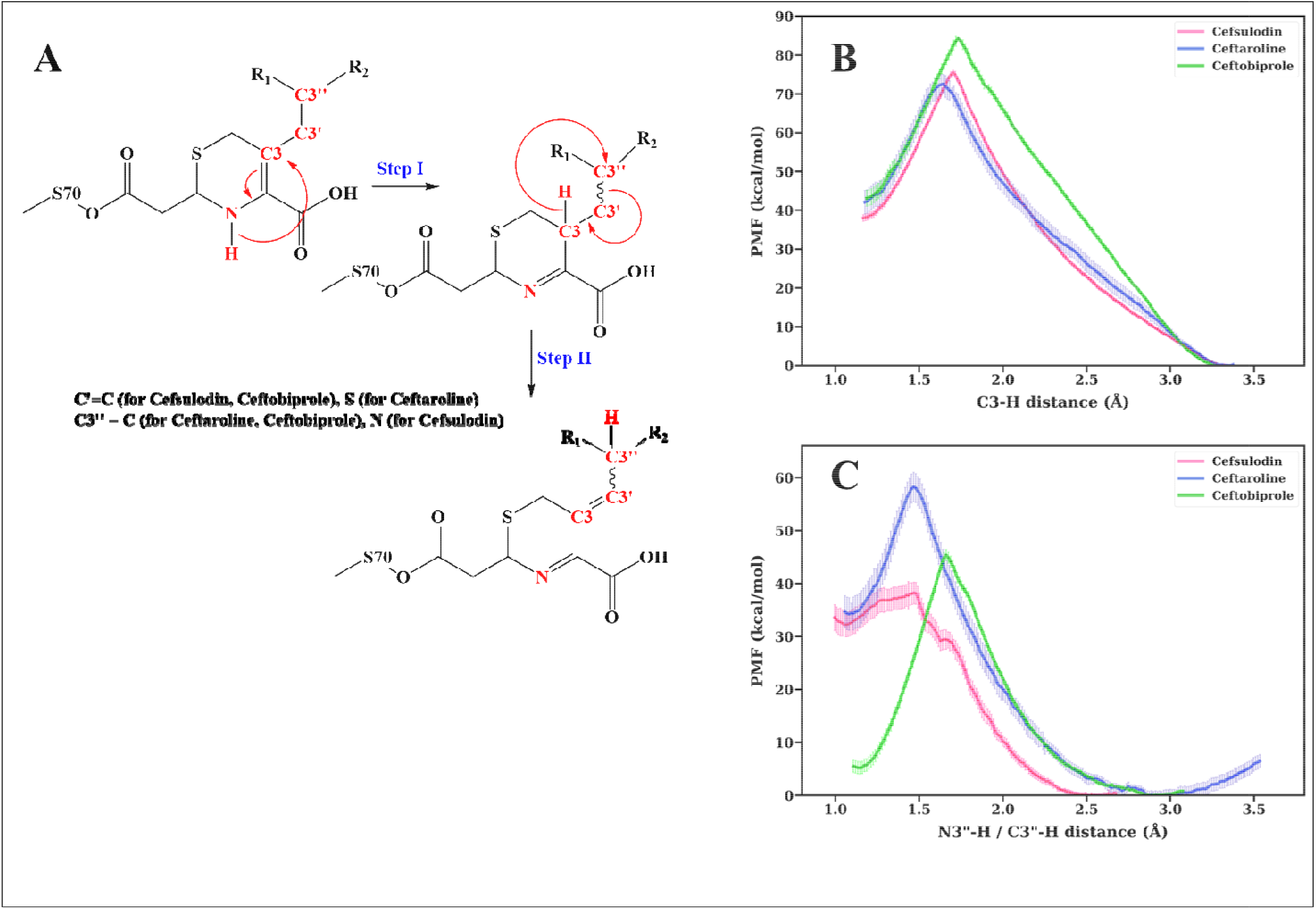
**(A)** The proposed two-step mechanism for the detachment of C3 sidechain, and the PMF values associated with **(B)** Step-I, and **(C)** Step-II. Here, and represents the side chains from Cefsulodin, Ceftaroline, and Ceftobiprole. The superscripts on the selected atoms are named for convenience. The dashed bond is kept intact to represent the diversification observed between Cefsulodin/Ceftaroline, and Ceftobiprole.

**Figure 9:**
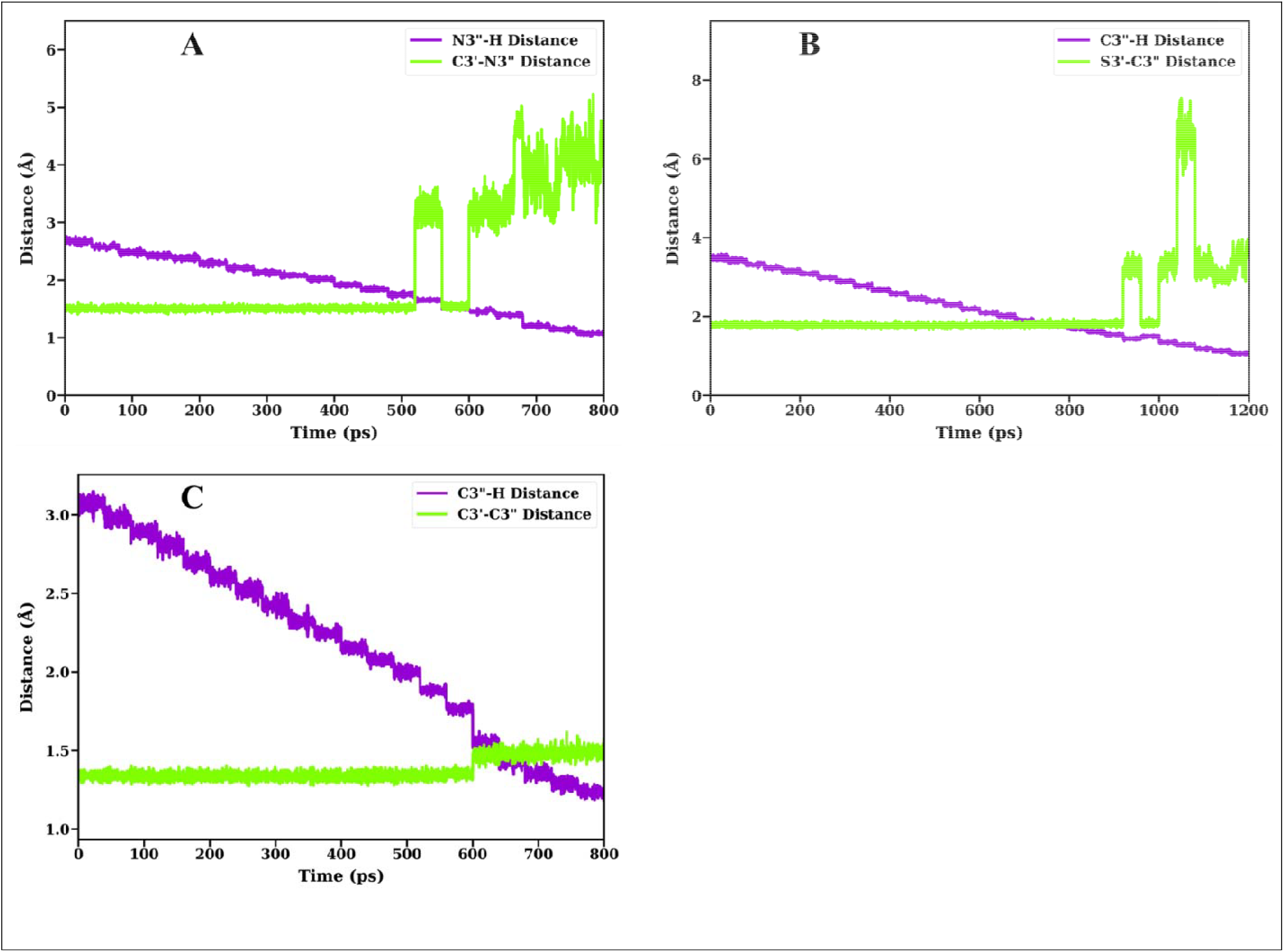
Distance variation plots for Step-II demonstrating C3 side chain detachment, as observed for. **(A)**Cefsulodin, **(B)** Ceftaroline, and **(C)** Ceftobiprole.

## Discussion

SME-1 can inactivate all classes of beta-lactam antibiotics, including 4th & 5th-generation cephalosporins with different C-3 substituents, making it an important target for AMR globally. There is a diverse variety in the C3 side chain of the cephalosporins, which allowed us to study the interaction of these molecules with SME-1. All the 5th-generation cephalosporins were poor substrates for the SME-1, which indicates that the bulkier C3 substitutions of cephalosporins directly desta ilize the binding of the cephalosporins to SME-1 Carbapenemase. Cefoperazone, with bulkier C7 substitutions than cefaclor, has a lower but yet significant value for catalytic efficiency with SME-1. Cefotetan with a unique dithietane ring in the C7 substituent also showed very poor turnover rates than Cefoperazone. These poorer turnover rates of catalysis are due to an additional H-bond of C-7 oxy-methyl substituent with Lys73 of the enzyme. Rest 4^th^ and 5^th^ generation cephalosporins showed very poor turnover rates that indicate resistance to SME-1 Carbapenemase. The efficiency of hydrolysis for the 4th and 5th generations was found to be much lower than that of older generations of cephalosporins, which can be attributed to the presence of bulkier C3 substituent. B-factor analysis of all the complex structures revealed that any cephalosporin does not stabilize the overall structure of the enzyme unlike the inhibitors of the enzymes. Notably, instead of stabilizing the complex, Cefoperazone and Ceftobiprole increased the overall b-factor of the enzyme. Indicating that bulky C7 side chains do not d any good for the enzyme’s stability. Our Mass spectrometric studies of SME E166A with these cephalosporins indicated that C7 side chains do not fragment on hydrolysis, but the C3 side chain does fragment in some cases only. The hydrolytic fragmentation of the C3 side chain depends on the C3’ atom and its hybridization state. Our detailed crystallographic studies, coupled with mass spectrometric analysis, revealed that the fragmentation of the side chain will take place where the C3’ atom is a carbon with sp3 hybridization. The bond between C3’ and C3’’ atoms will eventually break after hydrolysis, leaving an exomethylene group at the C3 position, the rest side chain will break from the cephalosporin. This fragmentation process seems to be independent of the C3’’ atom. We tested this diversity of the C3’’ atom with 3 different atoms like -O, -S & -N in cases of Cefotaxime, Cefotetan, and Cefpirome. The exact opposite of this fragmentation will occur when the C3’ atom is carbon but with sp2 hybridization. The double bond will undergo a shift, but the side chain will not fragment from the C3’ atom. We observed crystal structures of SME E166A with Ceftobiprole and Cefprozil to understand the mechanism and process for fragmentation. In both these complex crystal structures, the C3 side chain was still intact, indicating the crucial role of Hybridization in this hydrolytic fragmentation process. A similar trend was observed with the crystal structure of Ceftaroline Fosamil with SME E166A where C3’ atom is a sulfur. Sulfur having an empty d-block can behave differently than a sp3 carbon. The electron density for the side chain was quite clear, to say that due to the empty d-block of sulfur, the side chain of the Ceftaroline Fosamil remained intact upon hydrolysis. With cefovecin, the C3’ atom was part of a 5-membered ring, with the adjacent atom being an -O and an -C. So, it was expected to undergo fragmentation between C3’-O3’’ atoms. Indeed, the complex crystal structure of cefovecin showed the electron density of a fragmented linear side chain, which was completely different from the cyclic group substituent. It was predicted from the electron density that the C-O bond is more likely to be fragmented than the C-C bond, so the hydrolysed cefovecin was fitted as a C3’-O3’’ bond fragment. Our Mass spectrometric supported our crystallography data with Cefoperazone’s complex with SME E166A, clearly showing the mass difference for fragmented adduct. While Ceftaroline Fosamil, Ceftobiprole, and Cefixime showed unfragmented intermediate with SME-1 E166A.

X-ray crystallography coupled with Mass spectrometry gave us critical information regarding the hydrolysed and fragmented intermediates. But to confirm the mechanism of this C3 side chain fragmentation, we performed QM/MM simulations for the acyl-enzyme complexes of Ceftobiprole, Ceftaroline Fosamil, and Cefsulodin that represents our diverse sets of C3 side chain. A two-step proton transfer mechanism was proposed to explain the fragmentation of C3 side chain. The hydrolysed beta-lactam moiety contains a tertiary amine component, which partakes in an enamine-imine tautomerism, resulting in proton migration to the C3 atom(Figure 8(A)). Upon formation of imine, this C3 proton then moves to the C3’’/N3’’ atom, via formation of a transient 4-membered ring. If this second step occurs successfully, the side chain with a sp3 carbon at C3’ position will certainly break. But with a sp2 hybridized carbon, it only causes in a double bond shift from C3’’-C3’ to C3’-C3. With S at C3’ this proton shift could be difficult due to the vacant d-orbital over S, which might be involved in delocalization of π-electrons. The proton migration disrupts this extended delocalized π cloud, causing significant destabilization in the resulting system.

We calculated the PMF associated with both steps, and it was found that the energy required for the 1^st^ step was very similar across all three cephalosporins. Conversely, in 2^nd^ step, Cefsulodin demonstrated the lowest PMF, with the relative energy variations for Ceftobiprole and Ceftaroline standing at 6.82 kcal/mol and 20.05 kcal/mol, respectively. These observations indicate that the second phase might be the rate-limiting factor in this two-step process. This elucidates the C3 side separation noted for Cefsulodin but not for Ceftaroline. An intriguing observation is the reduced PMF value for Ceftobiprole in Step II, despite the absence of C3 side chain separation. The variation in distances between atom H-C3” and C3’-C3” was plotted to elucidate this phenomenon, as illustrated in Figure 9. In all the diagrams in Figure 9, the violet lines denote the migration of the H-atom from C3 to C3” (for Ceftaroline and Ceftobiprole), or from C3 to N3” (for Cefsulodin). A notable increase in the N3”-C3’/S3’-C3” distance is observed for Cefsulodin/Ceftaroline throughout this migration. This signifies the separation of the C3 side chain. The alteration in the C3’-C3” distance for Ceftobiprole was minimal, shifting from 1.3 to 1.5 Å, which accounts for the transition from a double bond to a single bond. Consequently, no dissociation of the C3 side chain was detected for Ceftobiprole.

## Conclusion

The comprehensive structural data and molecular simulations reported herein offer the most complete depiction to date of the hydrolytic fragmentation of the C3 side chain of cephalosporins by SME-1 Class A Carbapenemase. Our structural data shows the architecture of the active site of SME-1 Carbapenemase and how it accommodates the cephalosporins with diverse side chains. Our enzyme kinetics and ITC data complement the structural data and provide the key parameters of catalysis and the role of different C3 and C7 side chains in binding with the Carbapenemase. High-resolution crystallography structures reveal the nature of the C3 side chain while binding to the enzyme. It helps us establish the hydrolytic fragmentation of the C3 side chain, and C3’ atom’s nature and hybridization state determine whether the C3 side chain will fragment or not. QM/MM simulation further confirms our structural findings and provides the reasoning why the C3 side chain will be cleaved in sp3 C3’ carbon but not with sp2 carbon and sulfur. The variations in the distance of C3’-C3” in ceftobiprole from 1.3 to 1.5 Å confirmed the shifting of the pi cloud, but the fragmentation of the side chain does not take place. With Ceftaroline Fosamil, the presence of a vacant d-orbital over S atom might be involved in π-cloud delocalization, making it difficult for C3 side chain detachment.

## Materials and Methods

### Cloning, Overexpression & Purification

The SME-1 beta-lactamase gene from Serratia marcescens was cloned to get protein expression^16^. The chromosomal SME-1 gene sequence was first acquired from the Uniprot database. A plasmid based on pET-28a, including the SME-1 gene sequence, was utilized for further protein purification. This plasmid facilitates the expression of a C-terminal 6x His-tagged SME-1^17^ protein in Escherichia coli BL21/DE3 cells, with induction at 0.5 mM IPTG at an OD600 of 0.6. Following 18 hours of induction, cells were collected and resuspended in 50 mM Tris-HCl, 250 mM NaCl, pH 7.5, and subsequently disrupted via sonication with the incorporation of a protease inhibitor cocktail. The soluble fraction, derived from centrifugation, was filtered using a 0.45-μm syringe filter, applied to a Ni-NTA agarose column^18^, and eluted in succession with buffers comprising 55 mM Tris-HCl, 250 mM NaCl, pH 7.5, augmented with increasing concentrations of imidazole (10 mM and 250 mM). The eluted fractions from the 250 mM imidazole stage were concentrated utilizing an Amicon protein concentrator with a 10 kDa cutoff. Ultimately, protein purity was assessed utilizing SDS-PAGE. The Ni-NTA purified protein underwent additional purification via gel filtration utilizing the AKTA Start purification system equipped with a Superdex 75 pg column, which has a capacity of 120 ml. The protein was concentrated in the 10 KDa filter to a volume of 1 ml and thereafter injected into the column. Fractions for purified protein were obtained by analysing the chromatogram^19^.

### Enzyme Kinetics

Kinetic experiments were performed utilizing a Cary 60 UV spectrophotometer. Before initiating the kinetics, SME-1 underwent a buffer exchange with 50 mM PBS at a pH of 7.2. The antibiotics were solubilized in phosphate buffer and subsequently diluted to a working concentration of 1 mM^18,20,21^. The experiment entailed performing enzyme kinetics using a spectrum of optimal substrate concentrations ranging from 20 to 120 μM. The reaction was initiated by introducing SME-1 at 10-200 nM concentrations for all respective cephalosporins. The initial velocity (V0) was obtained from the absorbance versus time graph and plotted against substrate concentration to calculate enzyme kinetic parameters using the Michaelis-Menten equation. The turnover number was calculated by dividing the Vmax by the enzyme concentration in µM^22^.

### Isothermal Titration Calorimetry (ITC)

Isothermal titration calorimetry (ITC) for the SME_E166A deacylation deficient mutant was conducted using Malvern ITC-PEAQ equipment^23,24^. 50 µM protein was made in a phosphate buffer and introduced into the sample cell of the instrument, which has a capacity of 280 µL. All cephalosporins were formulated in the identical buffer as the protein, with concentrations varying from 400 µM to 700 µM. The cephalosporins solutions were introduced into a 40 µL syringe, and data acquisition commenced. Nineteen doses of 2 µL of cephalosporin were administered into the protein solution. All thermodynamic parameters were computed using the software supplied with the instrument. DMSO was used to make a stock solution for cephalosporins insoluble in aqueous buffer, and it was diluted in buffer with DMSO concentration kept at less than 5%. Identical concentrations of DMSO were used in the protein solution to minimize the heat of dilution^25,26^.

### Mass Spectrometry

All the mass experiments were performed in the XEVO G2-XS QTOF system with SME-1 deacylation deficient mutant to study the rearrangements in the cephalosporins structure bound with the enzyme. ESI Mass spectrometry was used to ionize the sample. SME-1 E166A with a concentration of 50μM was used with the cephalosporin’s concentration of 2mM with a final volume of 50μl. Samples were prepared in the acetonitrile-water system^27^.

### Crystallization

SME-1 and its E166A mutant were obtained and concentrated to 10 mg/ml. Ninety-six well-sitting drop plates (Art Robbins) were employed for preliminary screening experiments. The original screening experiment utilized PACT Premier, MIDAS Plus, LMB Screen (Molecular Dimensions), and Rigaku Wizard Screens. 0.8 µL of protein was combined with 0.8 µL of precipitant solutions from these screens on a 96-well plate containing a reservoir of 70 µL. The plates were sealed and maintained at a temperature of 20°C. Crystals were examined in diverse settings. Subsequently, manual precipitant solutions were formulated^28,29^, and crystallization was reiterated by Hanging Drop Plates from Molecular Dimensions, utilizing drop volumes between 1 and 3 µL. 20% PEG 4000 and 0.2M Lithium Chloride were chosen as the final conditions for both enzymes, utilizing 2 µL of enzyme and 2 µL of precipitant solution^30–32^.

### Data collection & refinement

Crystals of the SME-1 E166A mutant, acquired via the Hanging Drop method, were soaked in a solution of different cephalosporins. Soaking durations for all cephalosporins ranged from 5 to 30 minutes. A 15% Xylitol solution with the mother liquor was employed as the cryoprotectant following the soaking process. soaked crystals were subsequently mounted to the Goniometer base, and data was acquired using the Rigaku Micromax HF-007 apparatus equipped with a Hypix detector and a rotating anode (Cu: 1.54Å) source. A minimum exposure duration of 30 seconds was given, with the scan width maintained at under 0.25 degrees. The acquired diffraction data was further analysed using Crysalis Pro. The Aimless program in the CCP4 i2 package^33–35^ was utilized for scaling, and molecular replacement was conducted in CCP4 i2^36^ to obtain the initial model. Subsequent refining and hand reconstruction of the protein structure and ligand were conducted in WinCOOT. Following the successful refining and fitting of the ligand, the structures were submitted to the Protein Data Bank (PDB)^37,38^. The structure of SME-1 (9JNC) and its deacylation deficient mutant (9JND) was determined with the help of Protein X-ray crystallography. All the cephalosporins were soaked in the crystals of SME-1 E166A deacylation deficient mutant crystals. Different soaking times were given to different cephalosporins (Figure 1), and data were collected multiple times; only the best-fitted data are shown here.

### NMR Spectroscopy

^1^H & ^13^C NMR experiments were performed for the cefovecin, and cefovecin was treated with SME-1 to study the fragmentation pattern with the C3 side chain having tetrahydrofuran moiety. A solution of 20mg/ml cefovecin was prepared in 100% D2O with 50 mM sodium phosphate (tribasic) at pH 7.4^39,40^. For fragmented cefovecin a solution of 20mg/ml was prepared having 5μM SME-1 in 50 mM sodium phosphate (tribasic) at pH 7.4^41–45^. This solution was kept overnight at 4*C to allow the hydrolysis of the beta-lactam ring to complete. Then, the NMR spectra were acquired with the help of JEOL JNM-ECZR 500 MHz NMR with 500 scans for ^1^H and 2000 scans for ^13^C spectra^46^.

### Antimicrobial susceptibility testing

Antimicrobial susceptibility testing of SME-1 Carbapenemase was conducted on the top 10 E. coli strains. In the disk diffusion assay, the top 10 E. coli strains expressing SME-1 were cultivated in LB Broth until the optical density reached 0.6^47^. Bacteria were inoculated onto the MH Agar plates, and a disk containing respective cephalosporins^48^ was positioned on the surface of the agar. Plates were incubated at 37°C for 18 hours, and the zone of inhibition was quantified. The minimum inhibitory concentration (MIC) of the cephalosporins was assessed using microdilution 96-well plates containing Mueller-Hinton broth. The initial carbapenem concentration was 50 mcg/ml. Following the cultivation of the bacteria to an optical density of 0.6, it was diluted to achieve a concentration of 10^5^ CFU/ml, and 5 µl of the bacterial solution was dispensed into each well. Plates were incubated for 18 hours to determine the MIC value of the carbapenems^26^.

## Computational Methods

### Model Preparation

The initial structures for the three enzyme-inhibitor complexes were taken from experimental studies. All the ligands were covalently attached to Ser70 in its acetylated form due to the single point mutation E166A, which inhibited the de-acylation of hydrolysed β-lactam based ligands. The PDB structures of all the proteins were cleaned, and correct protonation states of the titratable residues at pH 7.4 were added via the H++ server^49^. The ligands were devoid of H atoms. Hence, all the ligand H’s were added, followed by the minimization of the entire ligand, along with the covalently attached Ser70 using GAFF^50^ in the Avogadro Modelling Package^51^. During this optimization, constraints were added to Ser70 atoms to avoid any change in their positions. For Cefsulodin, the pyridinium group in the C3 side chain was absent, which was also modelled and optimized using Avogadro. The backbone atoms of protein were modelled using ff14SB^52^ force field, with the ligand parameters generated using GAFF. The RESP charges on the ligand were generated with the AM1-BCC^53^ method, using the Antechamber^54^ module of Ambertools23^55^. The entire protein-ligand assembly was enclosed in a rectangular box of 14 Å buffer beyond any protein atom, which was solvated with TIP3P water molecules^56^. An appropriate number of ions were further added to neutralize the system.

### Classical Molecular Dynamics (CMD)

All three protein-ligand complexes were subjected to a similar pipeline of stepwise minimizations, using a combination of steepest descent and conjugate gradient methods^57,58^. This was followed by, strictly in the same sequence, NVT heating, NPT equilibration and 250 ns production run. The temperature was maintained at 310 K using Langevin Dynamics thermostat^59^. The pressure was stabilized at 1 bar using Berendsen barostat^60^, with a collision frequency of 1.0 ps^−1^. Restraints were applied on all the heavy atoms of the solute during equilibration, which were gradually lowered in each stage. The non-bonded cut-off was set to 10 Å, with the electrostatic interactions handled with the Particle Mesh Ewald (PME) method^61^. All the covalent hydrogen bonds were constrained to their equilibrium distances using SHAKE^62^.

### QM/MM studies: Steered Molecular Dynamics (SMD) and Umbrella Sampling (US) simulations

PM6 method was used to study the dynamics of the QM region^63^. The QM region consisted of all ligand atoms except the C7 side chain, which was cleaved at the Cβ-Cα bond of the carbonyl carbon. Additionally, the atoms of Ser70 up to the Cβ atom were included in the QM region. The C7 side chain is primarily not involved in the process of C3 detachment. Hence, its removal from the QM region significantly lowered the system size. An overview of the atoms in the QM region for all the systems are shown in Figure S8. It has been shown by *Catherine et* a*l*.,^13^ that upon binding with Ser70, imine-enamine tautomerism is exhibited by β-lactam based drugs, transitioning between Δ1, and Δ2 pyrroline components. As the exact mechanism for C3 side chain removal is not known, a two-step mechanism was proposed, wherein Step I involves the conversion of Δ2-enamine to Δ1-imine via migration of the H-atom^64^. In the second step, this H-atom moves toward the C3” atom, causing a possible detachment of C3 side chain from the rest of the molecule. A general map for this two-step process is outlined in Figure 8 (A).

To determine the energy change associated with this two-step process, Umbrella Sampling (US) studies were conducted^65^. The generation of frames for each of these steps was done using the Steered Molecular Dynamics (SMD) protocol^66^. In Step I, the initial frame for SMD was selected from the last frame of cMD run. Here, the collective variable (CV) was defined as the linear combination of N-H, and C3-H distances (Figure 8 (A)). For Step II of the process, the last frame of Step I was taken, with the CV being the combination of C3-H, and C3’’/N’’-H distances. Each SMD was performed for 200 ps, with a force constant of 200 kcal mol^−1^Å^−1^. During US studies, the CV was set as the C3-H distance for Step I, and C”/N”-H distance for Step II. A width of 0.1 Å was used, with the respective distance variations being 3.34 to 1.16 Å, 3.38 to 1.24 Å, and 3.28 to 1.18 Å, respectively, for Cefsulodin, Ceftaroline, and Ceftobiprole in Step I. For Step II, the corresponding variations were 2.68 to 0.99 Å, 3.54 to 1.23 Å, and 3.08 to 1.10 Å for the three ligands. Different force constants were required to achieve an effective overlap of the neighbouring windows. The values of the force constants for Step I and Step II are shown in Table S2 and S3, respectively, with their respective window-wise overlaps presented in Figure S9. In each US window, 10ps equilibration and 40ps production run was carried out. In both SMD and US studies, restraints were applied on all the atoms of the six-membered heterocycle, except N, H, and C with a force constant of 30 kcal mol^−1^ Å^−1^. This was done to prevent the cleavage of unwanted bonds during migration of H-atom. As the same atoms were restrained in each case, we expect the relative ordering of the PMF (potential of mean force) values to remain unchanged. For all QM/MM studies, switching-based cutoffs were implemented, with 8 Å and 10 Å being the lower and upper limits, respectively. As bonds were cleaved to define the QM domain, link atoms were added, with the atomic number of 1. Also, the SHAKE option was disabled for the QM region to achieve migration of H-atom.

## AUTHOR INFORMATION

### Author Contributions

The manuscript was written through the contributions of all authors. All authors have given approval to the final version of the manuscript. ‡These authors contributed equally.

## ACKNOWLEDGMENT

We would like to thank the Institute Instrumentation Centre, IIT Roorkee, and the Department of Bioscience and Bioengineering, IIT Roorkee, for providing research facilities. We would like to thank the Indian Institute of Technology, Ropar, for providing the Mass Spectrometry facility.

## ABBREVIATIONS

