## Supplementary Information for "Structural insights into the mechanism of C3 side chain fragmentation in cephalosporins with SME-1 class A carbapenemase"

**INDEX**

| **Figure/Table Number** | **Description** | **Pages** |
| --- | --- | --- |
| Table S1 | Data collection and refinement statistics for the crystal structures. | S1 |
| Table S2 | Window-wise force constants associated with Step-I of Cefsulodin, Ceftaroline, and Ceftobiprole | S2 |
| Table S3 | Window-wise force constants associated with Step-II of Cefsulodin, Ceftaroline, and Ceftobiprole | S2-S3 |
| Figure S1 | 2D interactions of Cephalosporins (1^st^-3^rd^ generation) with SME-1 E166A mutant | S3 |
| Figure S2 | 2D interactions of Cefpirome, Cefprozil, Ceftobiprole, Ceftaroline Fosamil with SME-1 E166A mutant. | S4 |
| Figure S3 | B-factor analysis of Cephalosporins with SME-1 E166A mutant. | S5 |
| Figure S4 | ^1^H-NMR of Cefovecin | S5 |
| Figure S5 | ^13^C-NMR of Cefovecin | S6 |
| Figure S6 | ^1^H-NMR of hydrolyzed and fragmented Cefovecin | S6 |
| Figure S7 | ^13^C-NMR of hydrolyzed and fragmented Cefovecin | S7 |
| Figure S8 | 2D, and 3D representations of the QM regions for Cefsulodin, Ceftaroline Fosamil, and Ceftobiprole. | S8 |
| Figure S9 | Window-wise overlap obtained from Umbrella Sampling studies for Step-I and Step-II of the QM/MM simulations for Cefsulodin, Ceftaroline Fosamil, and Ceftobiprole. | S9 |

| **Parameters** | **Cefaclor** | **Cefprozil** | **Cefotetan** | **Cefsulodin** | **Cefotaxime** | **Cefoperazone** | **Cefovecin** | **Cefpirome** | **Ceftobiprole** | **Ceftaroline** |
| --- | --- | --- | --- | --- | --- | --- | --- | --- | --- | --- |
| PDB ID | **9IOW** | **9UJ5** | **9UJA** | **9UJ6** | **9UJ9** | **9UJ7** | **9UJ8** | **9UJC** | **9UJB** | **9IV7** |
| Resolution range (Å) | **21.4-2.4** | **21.32-2.3** | **23.27-2.2** | **22.4-2.0** | **21.98-2.4** | **23.18-2.20** | **21.5-2.1** | **25.83-2.00** | **25.69-2.1** | **21.74-2.4** |
| Completeness (%) | 99.9 | 99.9 | 99.8 | 99.9 | 99.9 | 99.8 | 98.3 | 99.9 | 99.9 | 98.4 |
| Number of observations | **32640** | **86766** | **89674** | **159445** | **240992** | **188085** | **62189** | **115487** | **209787** | **69322** |
| Number unique | **8331** | **14976** | **10993** | **32693** | **27717** | **26555** | **12658** | **14812** | **28581** | **18330** |
| Final R, *R*_free_ | **0.17 0.26** | **0.18, 0.26** | **0.17, 0.25** | **0.2, 0.26** | **0.205, 0.27** | **0.16, 0.24** | **0.18, 0.25** | **0.16, 0.22** | **0.187, 0.23** | **0.19, 0.26** |
| Total number of atoms | **2126** | **4331** | **2123** | **4351** | **4234** | **4315** | **2170** | **4281** | **4362** | **3324** |
| Space group | **P 1 21 1** | **P 1 21 1** | **P 1 21 1** | **P 1 21 1** | **P 1 21 1** | **P 1 21 1** | **P 1 21 1** | **P 1 21 1** | **P 1 21 1** | **P 1 21 1** |
| Cell constants, a, b, c, *α*, *β*, *γ* | **35.3, 50.7, 61.91, 90, 98.6, 90** | **36.07, 51.24, 131.15 90, 93.88, 90** | **35.9, 50.5, 60.1, 90, 97.84, 90** | **36.2, 51.05, 131.32, 90, 92.78, 90** | **71.48, 51.5, 77.02, 90, 113.94, 90** | **71.59, 51.35, 78, 90, 114.05, 90** | **36.3, 60.6, 50.34, 90, 98.03, 90** | **36.52, 49.89, 61.15, 90, 99.3, 90** | **36.50, 51.36, 131.09, 90, 93.06, 90** | **35.62, 51.08, 129.74, 90, 90.12, 90** |
| Mean((I)/sd(I)) | **10.9** | **11.7** | **11** | **9.7** | **10.6** | **12.6** | **13.2** | **13.2** | **13** | **7.5** |
| Multiplicity | **3.9** | **5.8** | **8.2** | **4.9** | **8.7** | **7.1** | **7.8** | **7.8** | **7.3** | **3.8** |
| Average *B* factors (Å^2^) | **19.5** | **18.0** | **27** | **15** | **24** | **20** | **24** | **18** | **17** | **18** |

**Table S1:** Data collection and refinement statistics for the crystal structures.

| **Cefsulodin** | | **Ceftaroline** | | **Ceftobiprole** | |
| --- | --- | --- | --- | --- | --- |
| **C^(c)^-H^(b)^ distance (Å)** | **Force constant (kcal mol^-1^ Å^-1^)** | **C^(c)^-H^(b)^ distance (Å)** | **Force constant (kcal mol^-1^ Å^-1^)** | **C^(c)^-H^(b)^ distance (Å)** | **Force constant (kcal mol^-1^ Å^-1^)** |
| 3.34 | 400 | 3.38 | 400 | 3.28 | 400 |
| 3.25 | 400 | 3.29 | 400 | 3.20 | 400 |
| 3.15 | 400 | 3.19 | 400 | 3.10 | 400 |
| 3.08 | 400 | 3.09 | 700 | 3.00 | 400 |
| 2.97 | 400 | 2.89 | 900 | 2.92 | 400 |
| 2.87 | 400 | 2.78 | 1000 | 2.85 | 400 |
| 2.79 | 700 | 2.69 | 1000 | 2.76 | 600 |
| 2.66 | 700 | 2.58 | 1000 | 2.67 | 900 |
| 2.58 | 700 | 2.49 | 1000 | 2.58 | 900 |
| 2.47 | 700 | 2.39 | 1000 | 2.49 | 900 |
| 2.35 | 700 | 2.28 | 1000 | 2.39 | 900 |
| 2.25 | 700 | 2.21 | 1000 | 2.32 | 900 |
| 2.18 | 700 | 2.15 | 1000 | 2.25 | 900 |
| 2.08 | 700 | 2.06 | 1000 | 2.18 | 900 |
| 1.99 | 700 | 1.97 | 1000 | 2.13 | 900 |
| 1.90 | 700 | 1.87 | 1000 | 1.96 | 1200 |
| 1.82 | 900 | 1.79 | 1000 | 1.90 | 1200 |
| 1.75 | 1200 | 1.72 | 1800 | 1.82 | 2000 |
| 1.70 | 1800 | 1.68 | 1500 | 1.75 | 3000 |
| 1.68 | 1200 | 1.62 | 1900 | 1.72 | 3000 |
| 1.51 | 1000 | 1.59 | 1000 | 1.69 | 2000 |
| 1.42 | 1000 | 1.52 | 1000 | 1.65 | 2200 |
| 1.31 | 1000 | 1.44 | 1200 | 1.55 | 2000 |
| 1.22 | 1000 | 1.35 | 1200 | 1.45 | 2000 |
| 1.16 | 1000 | 1.24 | 1200 | 1.36 | 2000 |
|  |  |  |  | 1.27 | 1600 |
|  |  |  |  | 1.18 | 800 |

**Table S2:** Window-wise force constants associated with Step-I of Cefsulodin, Ceftaroline and Ceftobiprole.

| **Cefsulodin** | | **Ceftaroline** | | **Ceftobiprole** | |
| --- | --- | --- | --- | --- | --- |
| **N^(d)^-H^(b)^ distance (Å)** | **Force constant (kcal mol^-1^ Å^-1^)** | **C^(d)^-H^(b)^ distance (Å)** | **Force constant (kcal mol^-1^ Å^-1^)** | **C^(d)^-H^(b)^ distance (Å)** | **Force constant (kcal mol^-1^ Å^-1^)** |
| 2.68 | 400 | 3.54 | 400 | 3.08 | 600 |
| 2.59 | 700 | 3.45 | 600 | 2.98 | 600 |
| 2.48 | 700 | 3.34 | 900 | 2.89 | 600 |
| 2.42 | 700 | 3.26 | 1000 | 2.79 | 600 |
| 2.37 | 700 | 3.17 | 1000 | 2.69 | 600 |
| 2.27 | 900 | 3.09 | 1300 | 2.59 | 600 |
| 2.19 | 1000 | 2.99 | 1300 | 2.49 | 600 |
| 2.11 | 1200 | 2.89 | 1300 | 2.39 | 600 |
| 2.06 | 1300 | 2.79 | 1300 | 2.29 | 900 |
| 1.99 | 1300 | 2.67 | 1300 | 2.21 | 1100 |
| 1.89 | 1700 | 2.57 | 1300 | 2.11 | 1100 |
| 1.80 | 1500 | 2.44 | 900 | 2.03 | 1100 |
| 1.70 | 1500 | 2.36 | 900 | 1.94 | 1100 |
| 1.65 | 1300 | 2.28 | 1300 | 1.90 | 1600 |
| 1.58 | 1300 | 2.17 | 1500 | 1.85 | 2400 |
| 1.46 | 1500 | 2.07 | 1500 | 1.80 | 3000 |
| 1.46 | 1700 | 1.99 | 1500 | 1.74 | 2000 |
| 1.40 | 1500 | 1.88 | 1700 | 1.67 | 3000 |
| 1.35 | 1900 | 1.79 | 1700 | 1.66 | 2400 |
| 1.27 | 2100 | 1.69 | 1300 | 1.64 | 1200 |
| 1.22 | 1700 | 1.65 | 1900 | 1.55 | 800 |
| 1.16 | 1700 | 1.56 | 1900 | 1.46 | 800 |
| 1.08 | 1700 | 1.50 | 2600 | 1.37 | 800 |
| 0.99 | 1700 | 1.48 | 2700 | 1.29 | 800 |
|  |  | 1.46 | 2800 | 1.19 | 800 |
|  |  | 1.40 | 1900 | 1.10 | 800 |
|  |  | 1.33 | 1900 |  |  |
|  |  | 1.23 | 1300 |  |  |

**Table S3:** Window-wise force constants associated with Step-II of Cefsulodin, Ceftaroline and Ceftobiprole.

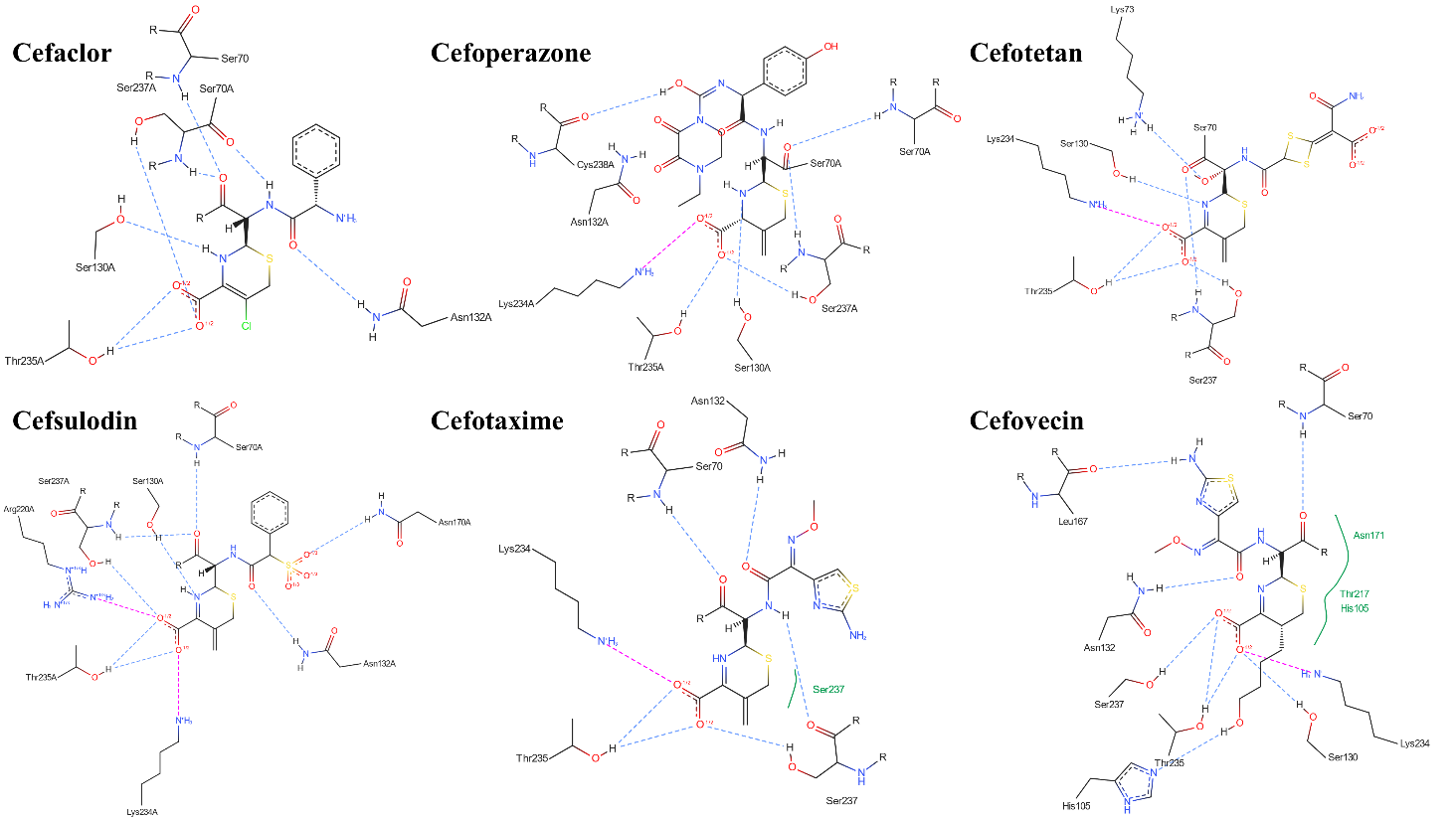

**Figure S1.** 2D Interactions of Cephalosporins (1^st^ -3^rd^ generation) with SME-1 E166A mutant.

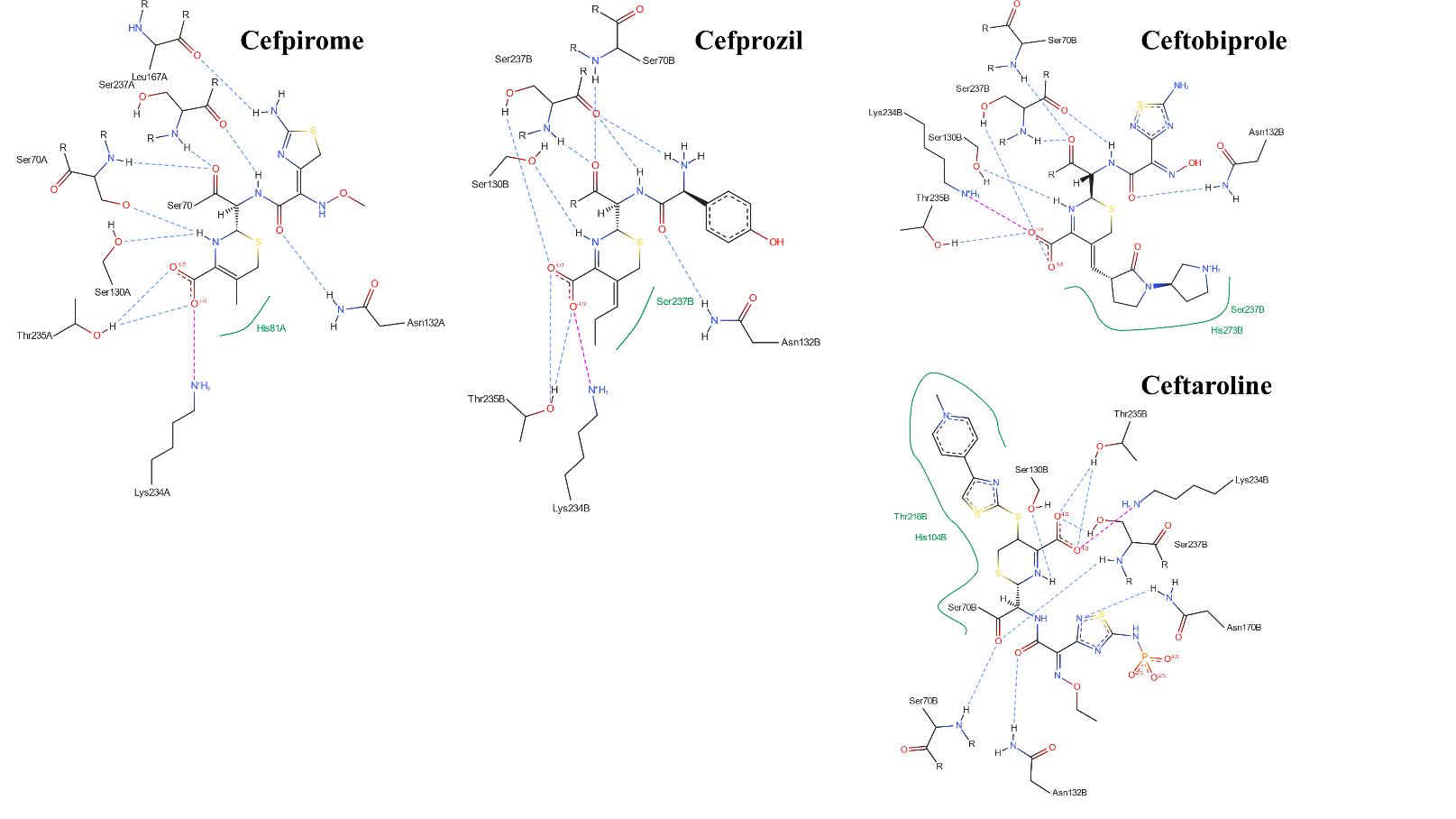

**Figure S2.** 2D Interactions of Cefpirome, Cefprozil, Ceftobiprole, Ceftaroline Fosamil with SME-1 E166A mutant.

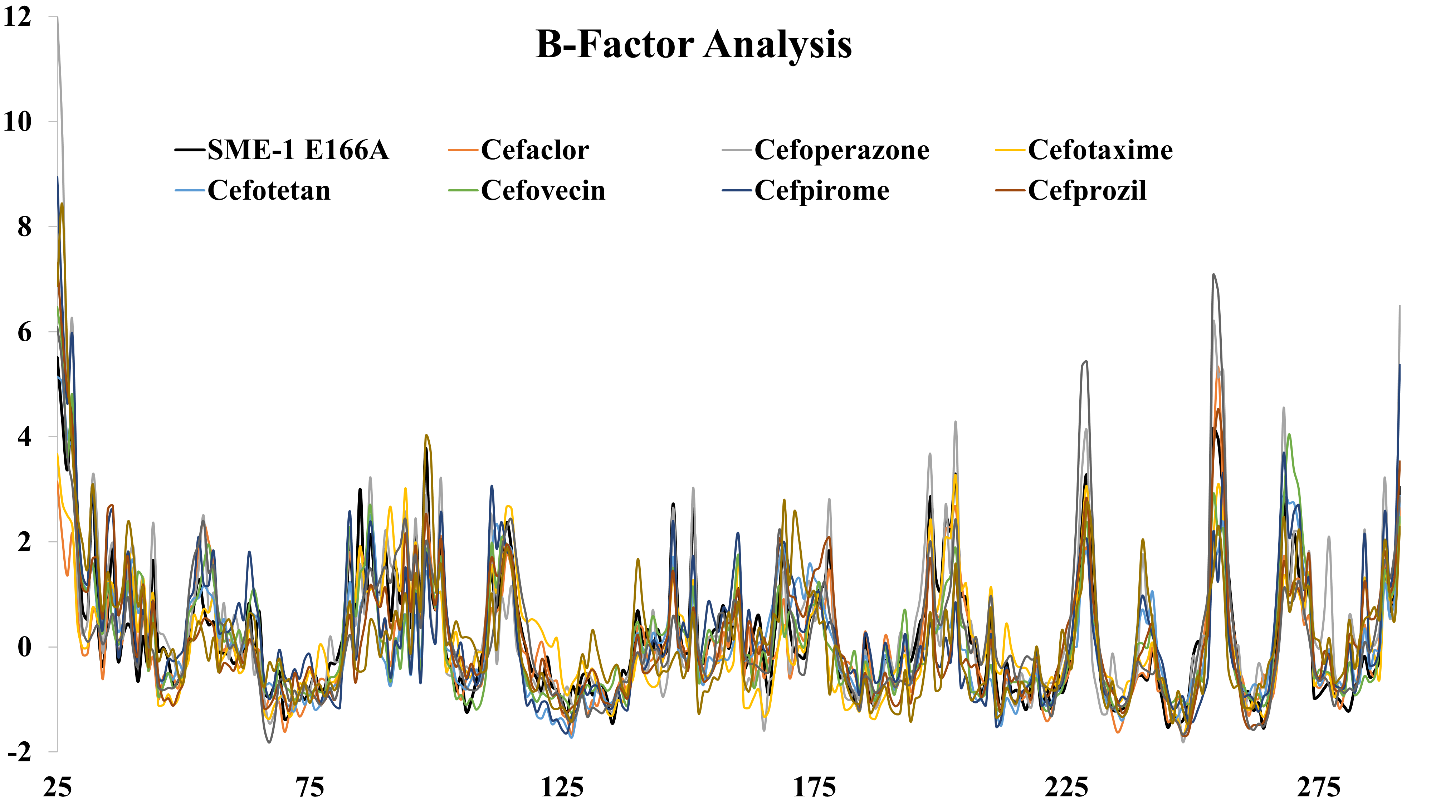

**Figure S3.** B-factor analysis of Cephalosporins with SME-1 E166A mutant.

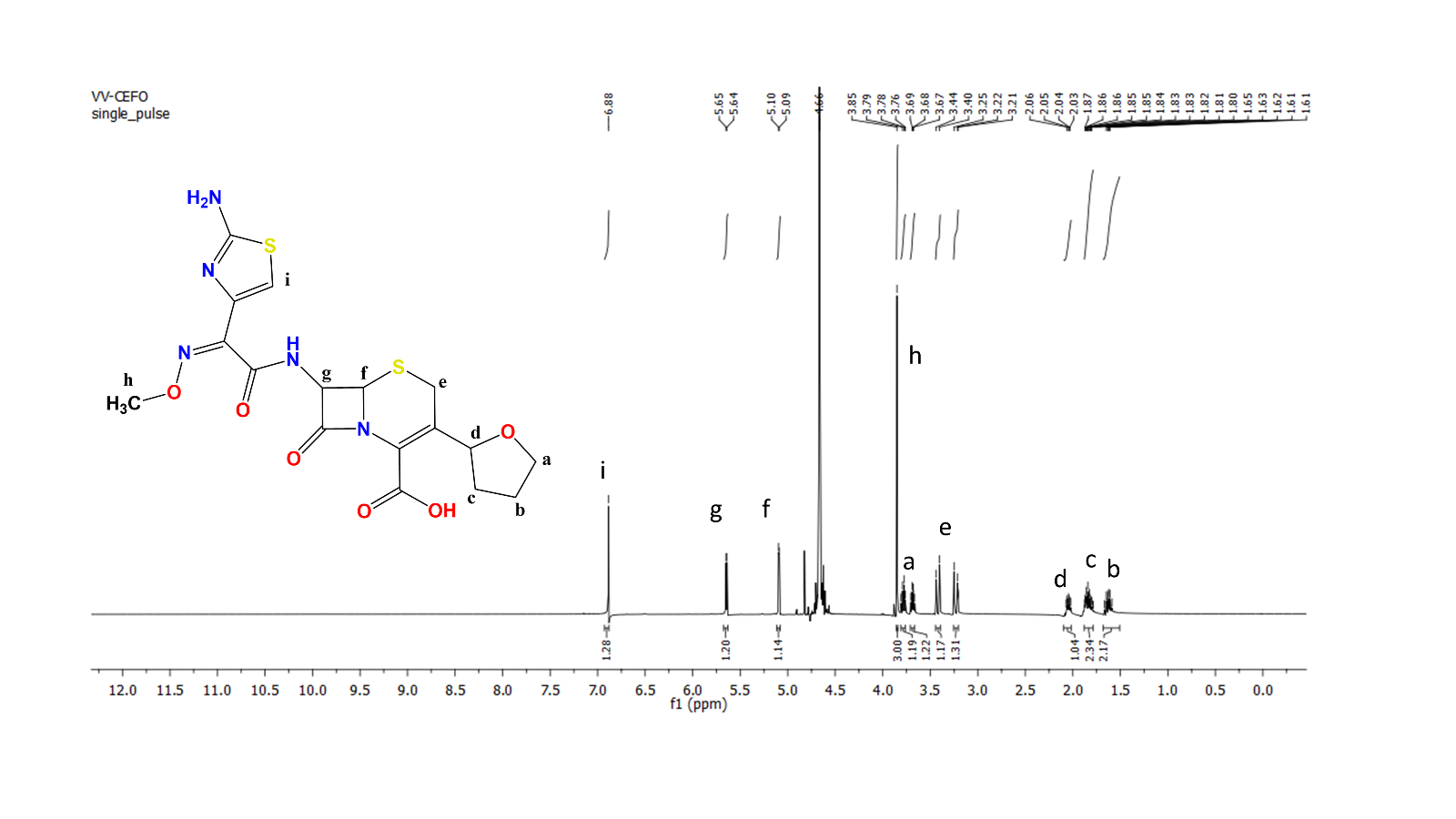

**Figure S4:** ^1^H NMR of Cefovecin.

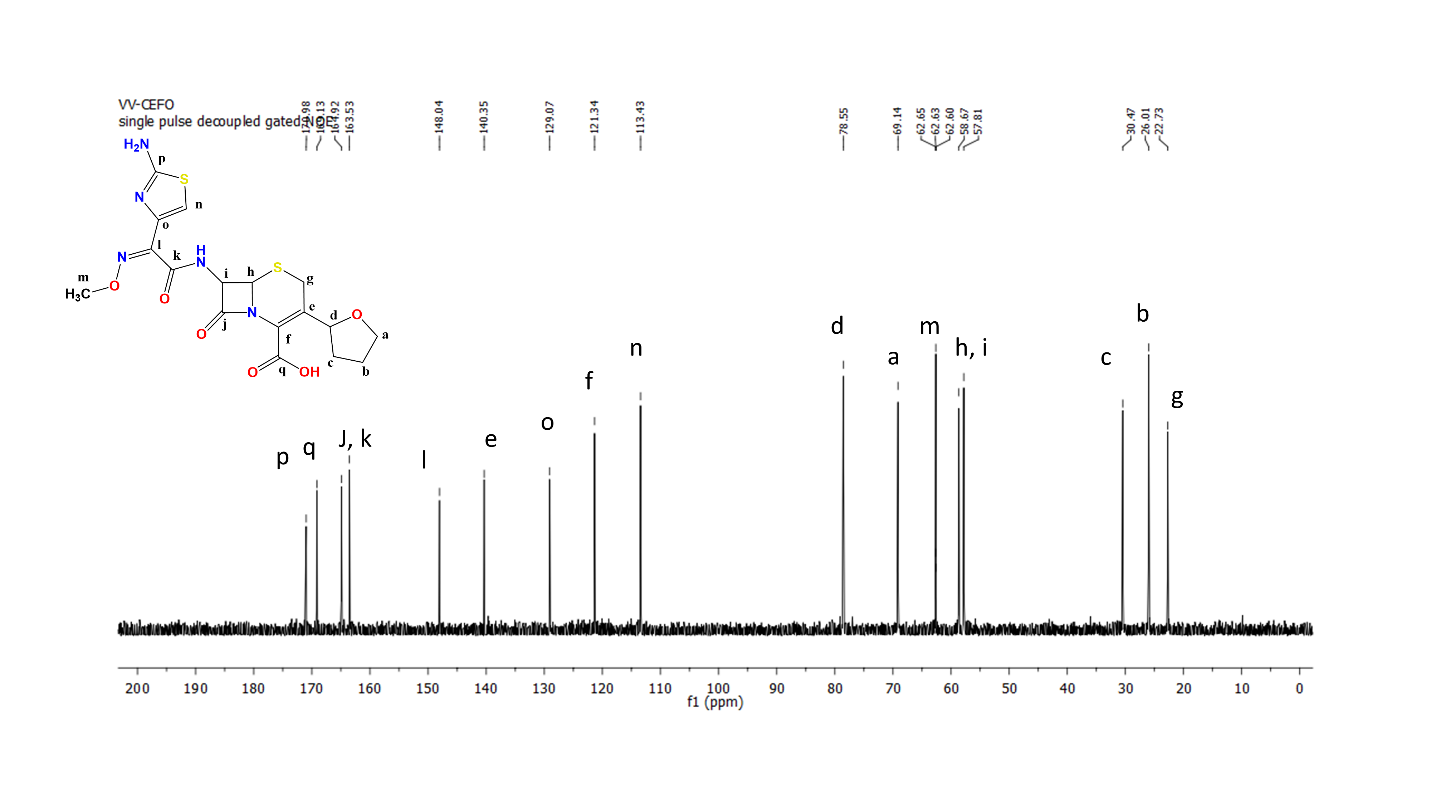

**Figure S5:** ^13^C NMR of Cefovecin.

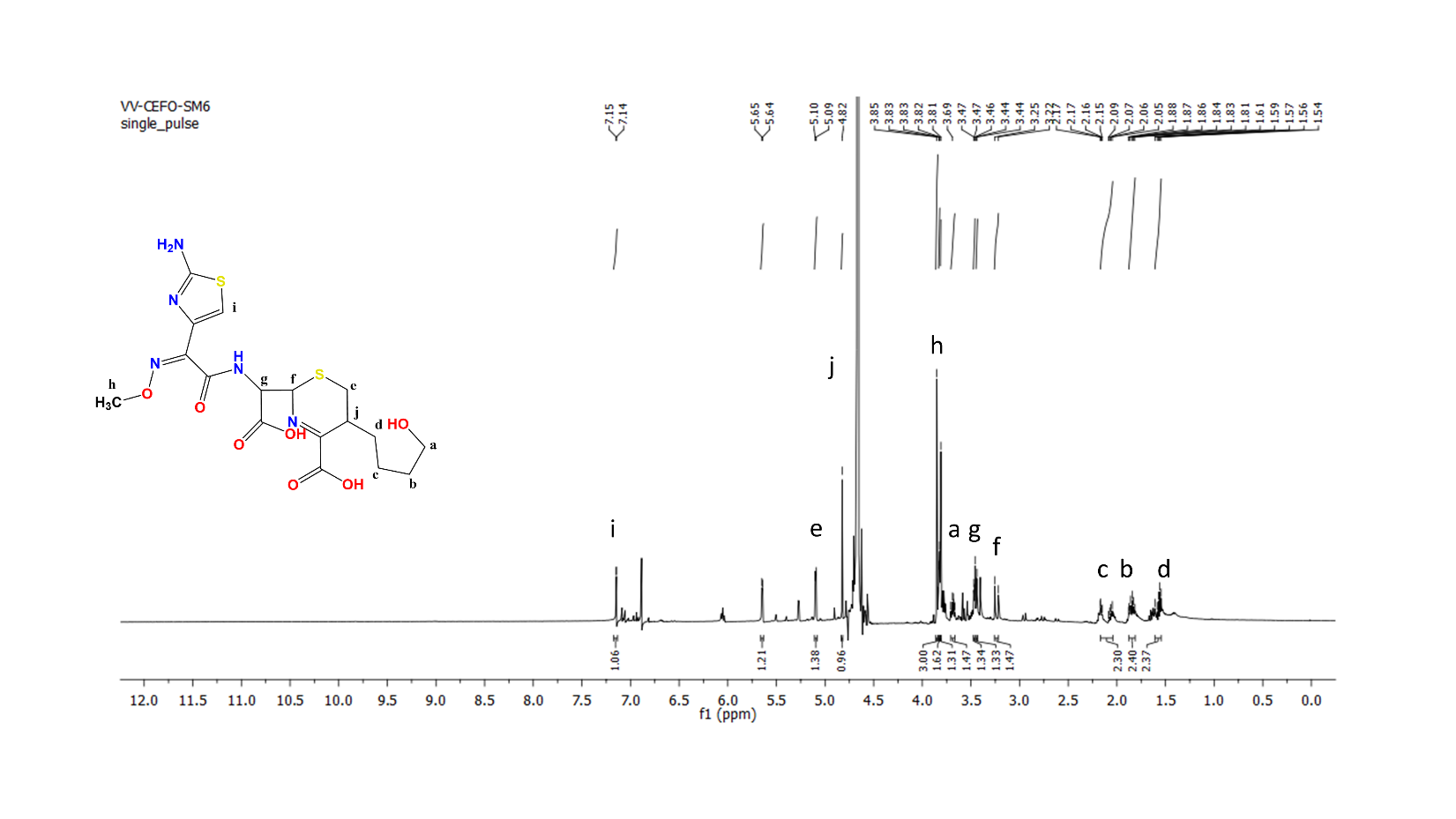

**Figure S6:** ^1^H NMR of hydrolyzed and fragmented Cefovecin.

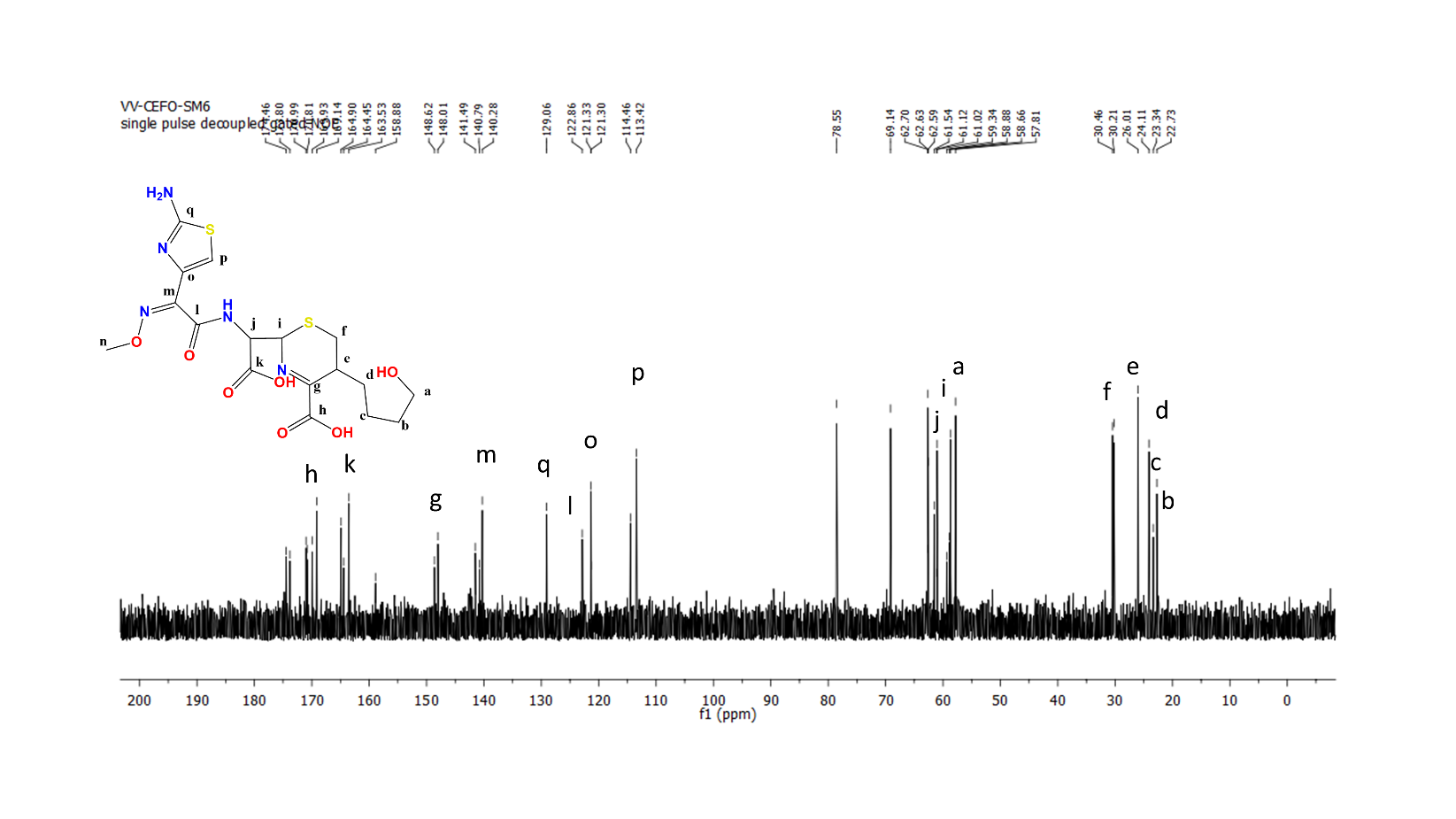

**Figure S7:** ^13^C NMR of hydrolyzed and fragmented Cefovecin.

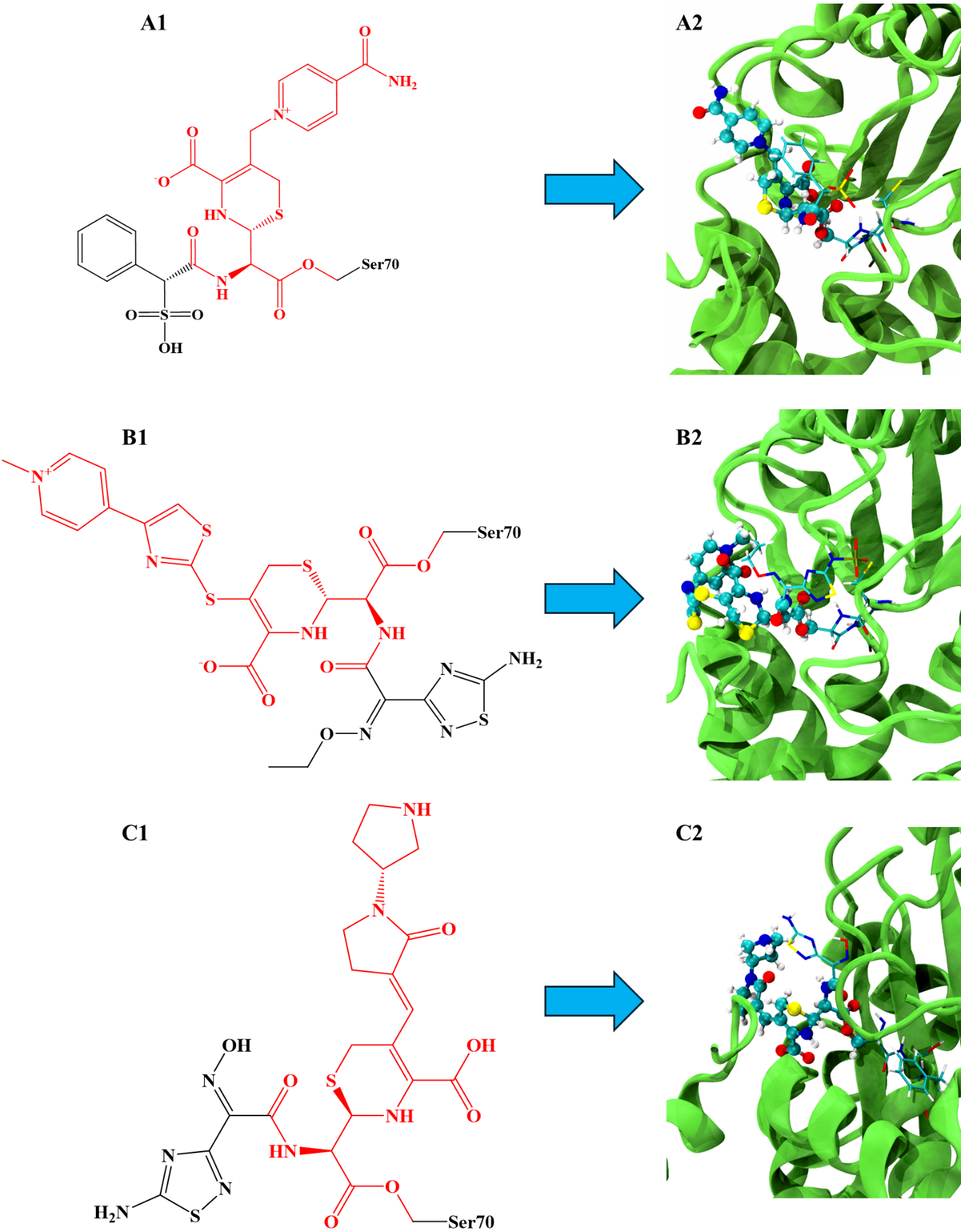

**Figure S8.** 2-D (Marked 1), and 3-D (Marked 2) representations of the QM region for Cefsulodin **(A)**, Ceftaroline Fosamil **(B)**, and Ceftobiprole **(C)**. The QM regions are highlighted in red (in 2-D representations); and with ball and stick model (in 3-D representations).

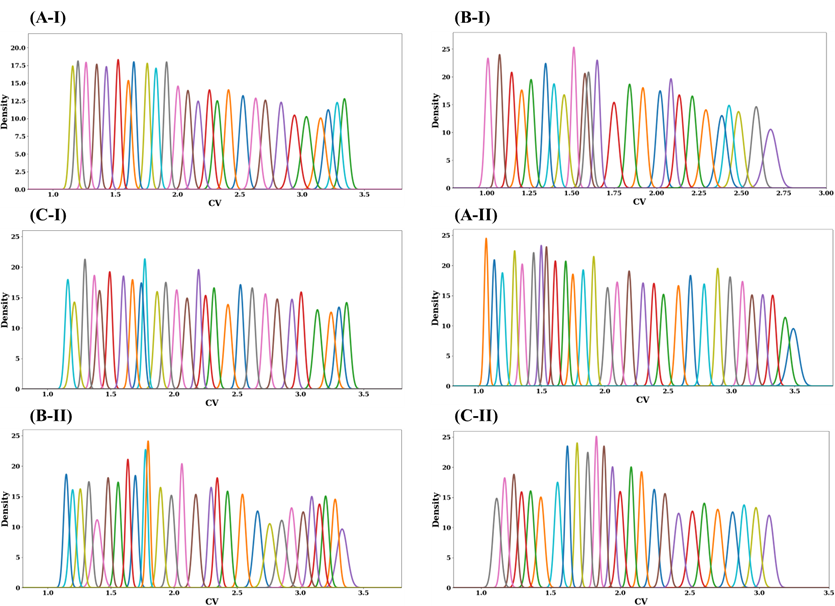

**Figure S9:** Window-wise overlap obtained from Umbrella Sampling (US) studies for Step-I (marked I) and Step -II (marked II), for Cefsulodin **(A)**, Ceftaroline Fosamil **(B)**, and Ceftobiprole **(C)**.
